# Naturalistic hyperscanning with wearable magnetoencephalography

**DOI:** 10.1101/2021.09.07.459124

**Authors:** Niall Holmes, Molly Rea, Ryan M. Hill, Elena Boto, Andrew Stuart, James Leggett, Lucy J. Edwards, Natalie Rhodes, Vishal Shah, James Osborne, T. Mark Fromhold, Paul Glover, P. Read Montague, Matthew J. Brookes, Richard Bowtell

## Abstract

The evolution of human cognitive function is reliant on complex social interactions which form the behavioural foundation of who we are. These social capacities are subject to dramatic change in disease and injury; yet their supporting neural substrates remain poorly understood. Hyperscanning employs functional neuroimaging to simultaneously assess brain activity in two individuals and offers the best means to understand the neural basis of social interaction. However, present technologies are limited, either by poor performance (low spatial/temporal precision) or unnatural scanning environment (claustrophobic scanners, with interactions via video). Here, we solve this problem by developing a new form of hyperscanning using wearable magnetoencephalography (MEG). This approach exploits quantum sensors for MEG signal detection, in combination with high-fidelity magnetic field control – afforded by a novel “matrix coil” system – to enable simultaneous scanning of two freely moving participants. We demonstrate our approach in a somatosensory task and an interactive ball game. Despite large and unpredictable subject motion, sensorimotor brain activity was delineated clearly in space and time, and correlation of the envelope of neuronal oscillations between people was demonstrated. In sum, unlike existing modalities, wearable-MEG combines high fidelity data acquisition and a naturalistic setting, which will facilitate a new generation of hyperscanning.

## 1. Introduction

Human social interaction is at the core of healthy neurodevelopment. From tactile stimulation to the evolution of language, from information transfer to social development, how we interact with others shapes everything from our abilities and skills to our personalities. However, relatively little is known about the neural underpinnings of these interactions. The simultaneous recording of functional brain imaging data from multiple people (hyperscanning) offers a powerful tool to probe brain activity underlying social interaction (Hari et al., 2015; Hari and Kujala, 2009). However, the available functional imaging technology places severe limitations on experimental design, participant experience, and data quality (Czeszumski et al., 2020). We aim to address these issues by developing a fundamentally new technique for hyperscanning, offering high performance, and the opportunity for naturalistic social interactions.

Functional magnetic resonance imaging (fMRI) offers assessment of brain activity with high spatial resolution, but the requirement that participants be enclosed and motionless in a noisy scanner makes natural interactions during hyperscanning impossible. Whilst MRI can be adapted to scan two people simultaneously (Lee et al., 2012; Renvall et al., 2020), this results in a claustrophobic environment which offers limited possibilities for experimental design. Most fMRI hyperscanning studies (King-Casas et al., 2005; Montague et al., 2002) have used separate scanners connected via video, but this also imposes barriers to natural interaction. In addition, the latency and longevity of the haemodynamic signal makes it challenging to assess brain dynamics.

In contrast, functional near infrared spectroscopy (fNIRS) (Ferrari and Quaresima, 2012) and electroencephalography (EEG) (Lopes da Silva, 2013) are wearable technologies that can be deployed in real-life settings (Dikker et al., 2017; Leong et al., 2017; Reindl et al., 2018), enabling more naturalistic hyperscanning. However, fNIRS suffers poor spatial resolution and (like fMRI) is limited to haemodynamic measurement. EEG, via assessment of scalp-level electrical potentials, directly measures brain electrophysiology and consequently has excellent temporal resolution, but suffers from poor spatial resolution and is sensitive to artefacts from non-neuronal sources of electrical activity, especially muscles during participant movement (Muthukumaraswamy, 2013).

Magnetoencephalography (MEG) measures the magnetic fields generated by neuronal currents (Cohen, 1968), providing direct assessment of electrophysiology. Unlike EEG, MEG has high spatial precision (Baillet, 2017; Hämäläinen et al., 1993) and lower sensitivity to non-neuronal artefact (Boto et al., 2019). However, MEG systems use cryogenically cooled superconducting quantum interference devices (SQUIDs) (Cohen, 1972) housed in magnetically shielded rooms (MSRs) to gain sufficient sensitivity to measure the neuromagnetic field. Low temperatures mean sensors are positioned in a fixed array, 2 – 3 cm from the scalp (to provide thermal insulation). So, like MRI scanners, MEG systems are cumbersome and static; only one person can be scanned at once, participants must remain motionless, and performance is limited by sensor proximity. Nevertheless, the potential for hyperscanning has been demonstrated using two MEG systems sited in the same MSR (Hirata et al., 2014), or geographically displaced systems connected via video (Baess et al., 2012; Zhdanov et al., 2015). Sequential dual-brain imaging studies have also been performed (Levy et al., 2021, 2017) with participants viewing videos of social interaction.

In sum, hyperscanning experiments can be carried out with existing technology, and such studies are beginning to provide unique insights into how the human brain mediates social interaction (Czeszumski et al., 2020). However, current instrumentation is limited either by its performance (EEG/fNIRS) or the unnatural scanning environment it provides (MEG/fMRI). The development of new technology which can scan two people during *live naturalistic interaction*, and provide *high spatiotemporal resolution, artefact free, data* could transform this field.

Recently, ‘wearable’ MEG has been developed through the use of optically pumped magnetometers (OPMs) (Boto et al., 2018). OPMs are sensitive magnetic field sensors that *do not require cryogenics*. These devices have enabled the design of flexible MEG sensor arrays which can be placed closer to the scalp, and adapted to the requirements of individual studies and participants. Increased proximity to the scalp improves sensitivity and spatial resolution beyond that which is achieved using cryogenic MEG (Boto et al., 2016; Iivanainen et al., 2017). In addition, the lightweight nature of OPMs has enabled the development of wearable systems which allow participants to move during recordings. This motion tolerance, coupled with provision of high-fidelity data, offers an ideal platform for hyperscanning.

However, to achieve sensitivity to the MEG signal, OPM-MEG requires a strict zero magnetic field environment (Allred et al., 2002). Further, OPMs are vector magnetometers meaning any movement of a sensor through a non-zero background field will generate artefacts that mask brain activity and can saturate sensor outputs. These constraints mean OPM-MEG experiments involving participant motion not only require an MSR, but also ‘active’ magnetic shielding in the form of electromagnetic coils which cancel any residual magnetic field experienced by the array (Borna et al., 2020; Holmes et al., 2019, 2018; Iivanainen et al., 2019). Such shielding systems have been shown to allow participant motion during MEG studies (Boto et al., 2018; Holmes et al., 2018). This has enabled the recording of brain activity in individual participants undertaking naturalistic tasks (Boto et al., 2018) and exploring virtual reality environments (Roberts et al., 2019), as well as the investigation of cognitive function (Tierney et al., 2018), cerebellar (Lin et al., 2019) and hippocampal (Barry et al., 2019; Tierney et al., 2021) activity, functional connectivity (Boto et al., 2021) and epilepsy (Vivekananda et al., 2020) using a lifespan compliant system (Hill et al., 2019). Such studies demonstrate the power of OPM-MEG as a neuroscientific tool. Nevertheless, active shielding has, until now, only enabled movements within a pre-specified volume at the centre of an MSR and present OPM-MEG systems could not be deployed for hyperscanning.

Here, we describe the technical developments required to overcome this limitation, thus allowing the first OPM-MEG hyperscanning experiments to be performed. The key enabling advance is the ‘matrix-coil’ system. Unlike previous active magnetic shielding systems, this allows accurate field control anywhere within the volume surrounded by the coil set. Moreover, by positioning two spatially separated zero-field regions over OPM arrays worn by interacting participants, we provide the environment needed for the collection of high-quality MEG data in two-person experiments. In what follows, we describe the matrix coil and provide examples of its use in both hyperscanning and single-participant experiments.

## 2. Results

### Two-person touching task

We first explored the capabilities of OPM-MEG hyperscanning by conducting a simple, guided, two-person touch experiment. Each participant wore an OPM-MEG helmet containing 16 OPMs placed over left sensorimotor cortex. Each OPM is a small integrated unit incorporating a heated cell containing a vapor of rubidium-87 atoms, a 795-nm wavelength diode laser tuned to the D1 transition of rubidium, and a photodetector. Following optical pumping, the rubidium atoms are insensitive to photons of the polarised laser light at zero magnetic field and thus the intensity of the light which passes through the cell to the photodetector is at a maximum. Changes in the magnetic field experienced by the cell result in a decrease in the measured laser intensity as photons are absorbed by the atoms (Dupont-Roc et al., 1969; Shah and Wakai, 2013). This means the photodetector signal can be used as a sensitive measure of magnetic field. Our OPM-MEG system uses arrays of QuSpin Inc. (Louisville, Colorado, USA) Zero Field Magnetometers, which each have a dynamic range of ±5 nT, a noise-floor of <10 fT/√Hz, and a bandwidth of 0-130 Hz (Shah et al., 2018). The OPMs were located on the scalp using 3D printed helmets and a co-registration procedure (Boto et al., 2017; Hill et al., 2020) provided information about the sensor locations relative to brain anatomy.

The experimental setup is shown in Figure 1a. Two participants stood either side of a table, ∼65 cm apart. The matrix coil was used to null remnant magnetic field inside the MSR, at the locations of both helmets (thus allowing natural movement for both participants). Upon hearing an audio cue, participant 1 (female, right-handed, age 30, height 172 cm) reached over the table with their right hand and stroked the back of the right hand of participant 2 (male, right-handed, age 25, height 182 cm). Upon a second (different) audio cue, the roles were reversed. Trials were defined as either ‘odd’ (participant 1 touches participant 2) or ‘even’ (participant 2 touches participant 1). The sequence was repeated 30 times (60 trials total) with an inter-trial interval of 5 seconds. The movements of the two OPM-MEG helmets were tracked using an optical tracking system.

**Figure 1:**
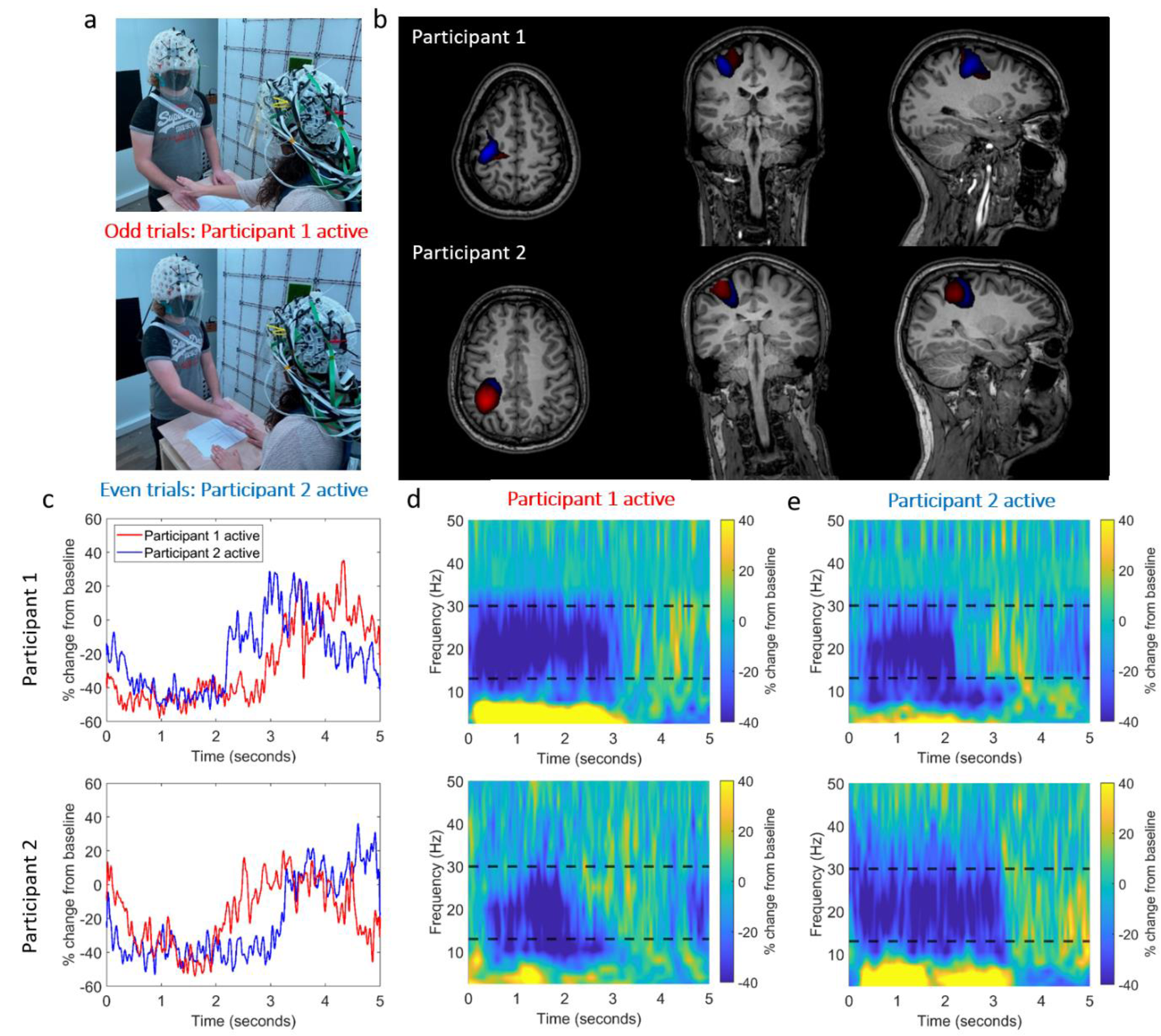
OPM-MEG data collected during a two-person naturalistic touching experiment. a) Two participants stood either side of a table. In odd numbered trials, participant 1 strokes the right hand of participant 2, with their right hand. In even numbered trials, the roles are reversed. b) Beamformer images show the spatial signature of beta band modulation (thresholded to 80% of the maximum value). The odd trials (participant 1 active) are shown in red and the even trials (participant 2 active) are shown in blue. The spatial pattern suggests activity in the sensorimotor regions. (Note there is a large overlap so the blue overlay is partially obscured). c) Timecourse showing the trial averaged envelope of beta oscillations, extracted from left sensorimotor cortex (peak in the beamformer images). Data from participants 1 and 2 are shown in the top and bottom rows respectively. In both cases, the red trace shows data recorded when participant 1 was touching participant 2. The blue trace shows data recorded when participant 2 was touching participant 1. d) Time-frequency spectra of activity in left sensorimotor cortex. Data were recorded when participant 1 was active. Upper plot shows data from participant 1, lower plot shows data from participant 2. The black dashed lines show the beta band. e) Equivalent to (d) but data shown for when participant 2 was active. Photographs are of the authors.

We hypothesised that beta band (13 – 30 Hz) modulation, as a result of the motor control or sensory response, would be observable in the primary sensorimotor regions. To test this, a beamformer approach (Vrba and Robinson, 2001) tuned to the beta band, was used to derive images of oscillatory modulation during the task. We also performed time-frequency analysis to show modulation of neural oscillations at the location of maximum beta modulation. Figure 1b shows beamformer images contrasting the task (0.5 s < t < 2 s) and control (3 s < t < 4.5 s) time windows. Figures 1c-e show the temporal dynamics of oscillatory power at the peak voxel location. Despite the large head movements which participants made as they reached across the table (maximum translations from the starting position in any one trial were 16 mm and 24 mm, for participants 1 and 2 respectively, the maximum rotations were 3.0° and 7.9° - see supplementary material) the expected task induced reduction in beta amplitude was observed. In each case, the active participant (i.e. the one performing the touch) showed a reduction in beta power that commenced earlier and persisted longer than that seen in the passive participant. This experiment demonstrates that high-quality OPM-MEG hyperscanning data can be obtained using our system, even in the presence of movements.

### Two-person ball game

To further demonstrate the system’s capabilities, we aimed to show that OPM-MEG hyperscanning can be used to measure brain activity whilst two players hit a table-tennis ball back and forth to one another. Unlike our guided touch task, which where we expected temporally smooth head movements, we expected this task to generate movements that were quicker and more unpredictable. Despite such movement (maximum translations from the starting position in any one trial were 50.0 mm and 64.8 mm, for participants 1 and 2 respectively, the maximum rotations were 17.0° and 17.4°) we hypothesised that the matrix coil system would reduce the remnant field sufficiently to enable collection of useful data; we expected to observe a decrease in beta power in left motor cortices during the task.

The experimental setup is shown in Figure 2a. Participant 1 (see above) and Participant 3 (male, right-handed, age 41, height 188 cm) undertook this experiment. The participants stood ∼80 cm apart, each holding a table-tennis bat in their right hand. The remnant magnetic field was nulled using the matrix coil system at the locations of both helmets. An audio cue signalled the participants to begin playing the game, after 5 seconds a second cue signalled the participants to stop the rally and rest for 7 seconds. This process was repeated 25 times and movement of the helmets during the experiment was recorded. Data were processed using a beamformer to derive an image showing the spatial signature of beta modulation between task (2 s < t < 4 s) and control (10 s < t < 12 s) windows. A time frequency spectrum was also extracted from the peak of the beamformer image. Figure 2b shows the beamformer images overlaid on an anatomical MRI. The spatial signature suggests activation in the motor cortex as expected. Figure 2c shows the time-frequency dynamics of oscillatory power, revealing a reduction in the amplitude of beta activity during the task. In addition to overall beta modulation (i.e. the difference between playing the game and resting) we also expected that, following each strike of the ball, a small amplitude increase in beta power should occur; assuming consistent timings, we expected this effect should alternate between participants (e.g. we expect a peak in activity for participant 1, and a trough in participant 2 just after participant 1 has hit the ball). Analysis was performed to probe the presence of this relationship. Figure 2d shows beta envelopes from both participants; data in the 3 s to 6.5 s time window were extracted and are shown inset in Figure 2e (blue for participant 1, red for participant 2). Autocorrelations of the two timecourses were computed and compared with their cross-correlation (Figure 2e). These data reveal the beta envelopes evolve in anti-correlation, with a lag of ∼0.6 s between participants. This direct observation of the correlation of the amplitude envelope of oscillatory brain activity in two participants carrying out a single task highlights the power of hyperscanning.

**Figure 2:**
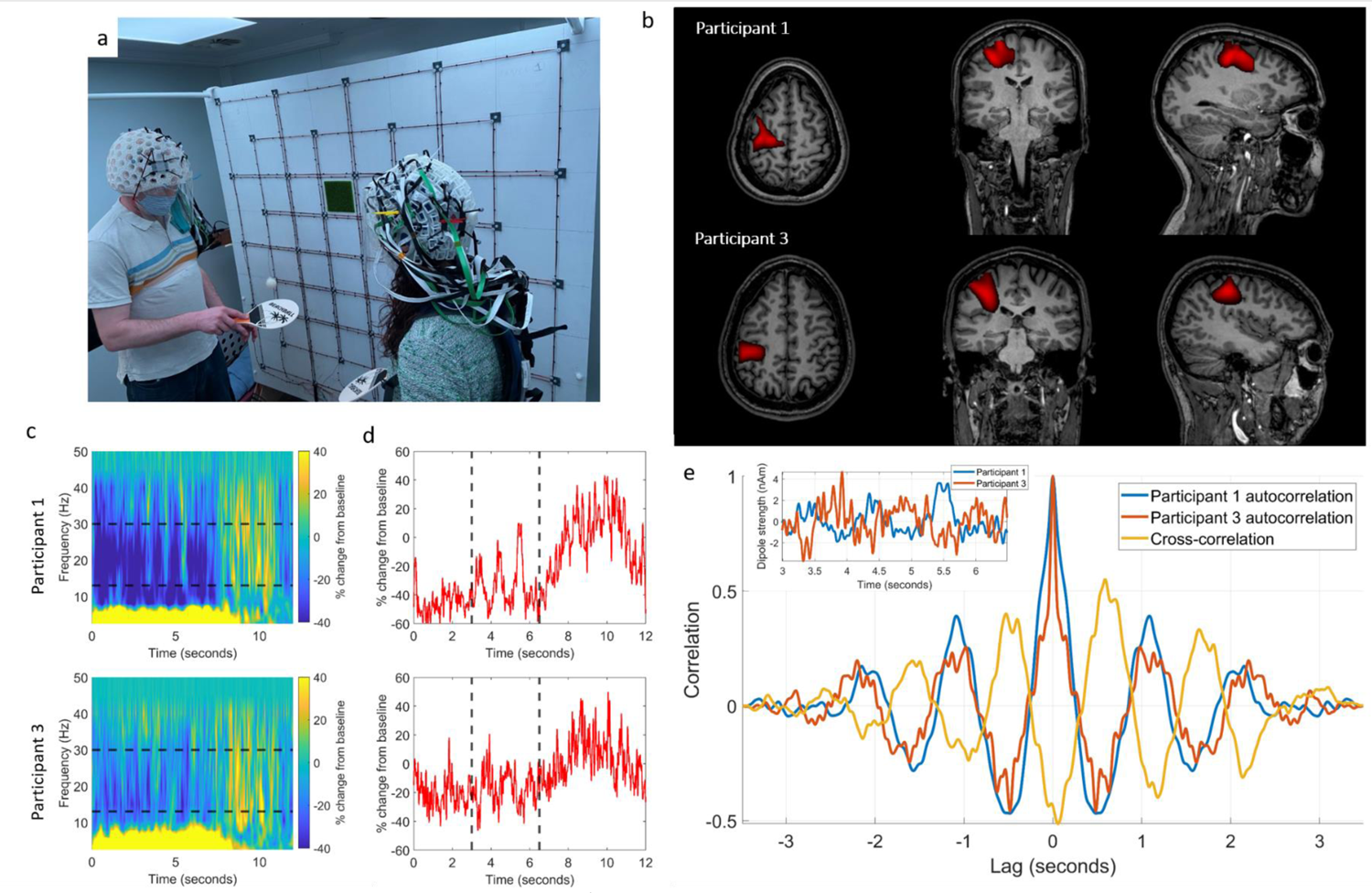
**OPM-MEG data collected during a two-person ball game**. a) Two participants each held a table-tennis bat in their right hand and hit the ball back and forth to each other; a 5 s rally was followed by 7 s rest. b) Beamformer images of beta band modulation between task and rest (thresholded to 80% of the maximum value). The spatial pattern suggests beta power reduction in the sensorimotor regions during the rally. c) Time-frequency spectrograms, extracted from left sensorimotor cortex, for participant 1 (top) and participant 3 (bottom). Black dashed lines show the beta band. d) Timecourses of the envelope of beta band activity (again participant 1 top, and participant 3 bottom). Data suggest anti-correlation between 3 and 6.5 seconds (marked with black dashed lines). e) Inset: the timecourses extracted in the 3 – 6.5 second window and overlaid. Main: comparison of the autocorrelations of the two extracted timecourses (blue/red) with their cross-correlation (yellow) reveals anticorrelation with a lag of ∼0.6 s between the participants’ brain activity. Photographs are of the authors.

### Matrix coils

OPM-MEG hyperscanning was made possible by our matrix coil, which acts to reduce the strength and spatial variation of the magnetic field surrounding the OPM arrays. This is critical since, to obtain the sensitivity required for MEG, OPMs must be operated at zero-field (Allred et al., 2002; Happer and Tam, 1977; Savukov and Romalis, 2005) requiring sensors to be screened from all sources of static and dynamic magnetic fields. This is achieved, in part, by operating inside MSRs constructed from multiple layers of high magnetic permeability material (mu-metal). However, the presence of the mu-metal leaves a remnant field which can be several 10’s of nanotesla (Boto et al., 2018; Hämäläinen et al., 1993; Holmes et al., 2018), meaning that active compensation using electromagnetic coils, which generate an equal and opposite magnetic field to that experienced by the array, is necessary.

OPMs typically feature ‘on-sensor’ coils, which can compensate local static magnetic fields experienced by the OPM, up to ±50 nT. Data are then measured relative to this offset within a narrow dynamic range of around ±5 nT. However, since this compensating field is set at the start of a MEG recording, any subsequent movement of an OPM with respect to the background field induces a change in the measured magnetic field, which affects the data in three main ways:

1. In the worst case, a field shift >5 nT will saturate sensor outputs so that no data can be collected (Holmes et al., 2018; Iivanainen et al., 2019).
2. Even if field shifts are smaller than the dynamic range, the accuracy of measured data is affected by a change in sensor gain; such changes can be as large as 5% for a 1.5 nT field offset (Boto et al., 2018); this nonlinearity causes a significant degradation of data fidelity.
3. Even in cases where a change in field does not cause appreciable modulation of sensor gain, the artefacts caused by rotating the sensor in a field, or translating it in a field gradient, can mask brain activity (e.g. in a 5 nT field, a rotation of just 1° would cause an artefact of ∼90 pT, which is ∼100 times larger than a typical evoked signal at the scalp (Boto et al., 2017)).

For these reasons, creating a zero magnetic field environment is crucial for enabling OPM operation.

To address these issues, we developed a system of matrix coils, comprising two 1.6 x 1.6 m^2^ planes, each containing 24, individually controllable, square coils. Each coil has a square side length of 38 cm and is formed of 10 turns of copper wire. A regular 4 x 4 grid of these coils is wound onto each plane along with an overlapping (to allow finer field control in off-axis directions) 3 x 3 grid of coils (excluding the central coil), as shown in Figure 3a. The coil planes were separated by 150 cm and positioned such that the centre of the coil array was at a height of 130 cm from the floor, with the array spanning a height range of 50 to 210 cm. By measuring the remnant magnetic field inside the MSR experienced by each OPM (projected along its sensitive axes) in the helmet, along with a calibration matrix containing the magnetic field generated per unit current, at each OPM, by each of the 48 matrix coils, we can compute the coil currents that will optimally null the magnetic field experienced by the array. This data-driven approach is easily extended to two separate arrays, and provides a low field environment for OPM-MEG experiments which can be readily adapted to the requirements of different paradigms. Participants are required to remain still during the nulling process, but the nulled volumes can be placed at any location between the coils, meaning that an experiment can be carried-out with a single participant standing or seated, or with multiple participants.

**Figure 3:**
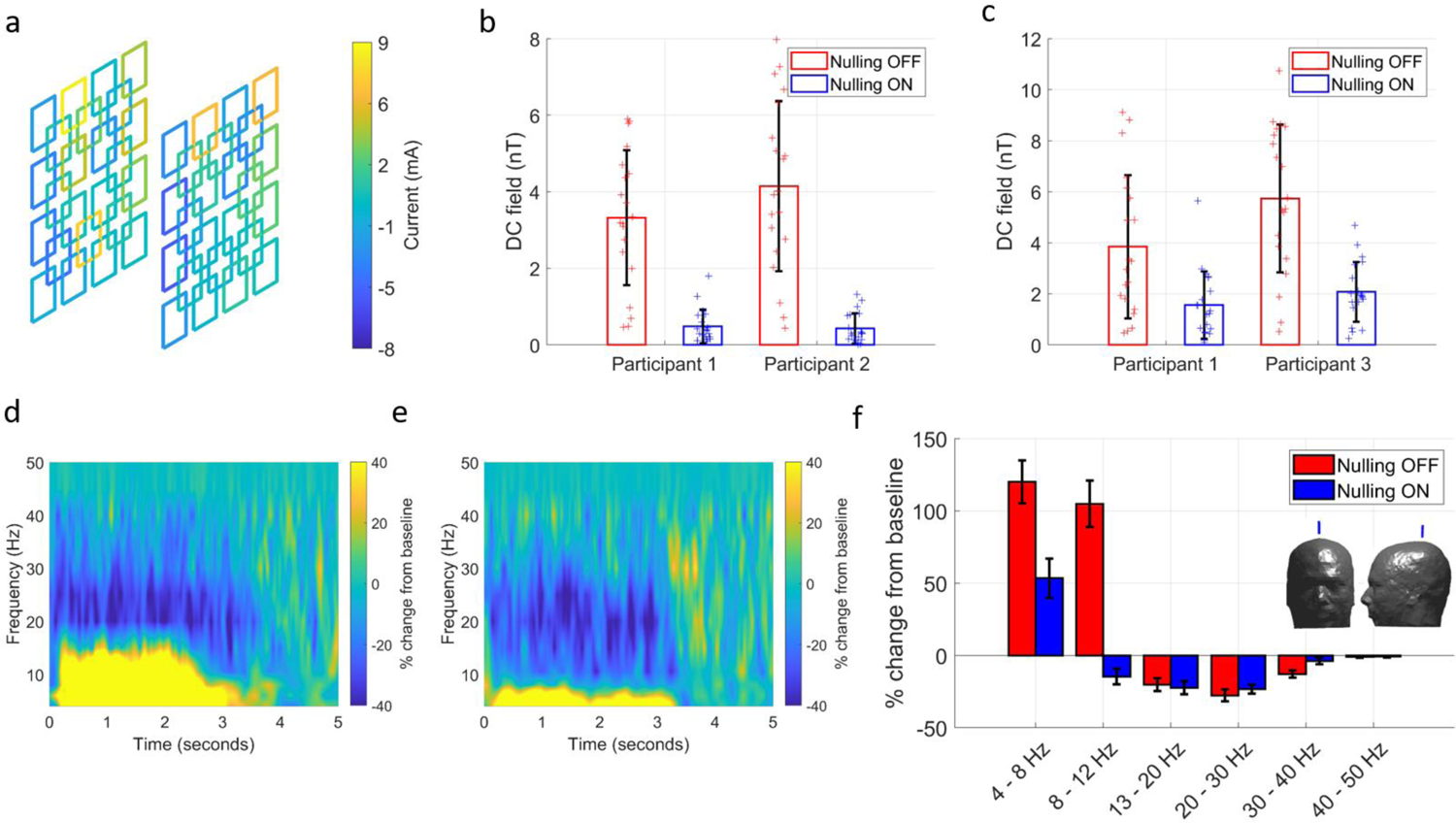
The matrix coil system and its effects on the background magnetic field and data quality. a) The 48-coil bi-planar matrix coil system. The current in each coil is individually controlled to generate the required field in order to cancel the remnant magnetic field inside the MSR. The current distribution shown was used during the two-person ball game. b) The strength of the DC field, reported by the 48 total field measurements from 24 OPMs (12 per participant), with and without the matrix coils active, during the two-person touch task. The error bars show standard error across sensors. c) Equivalent to (b) for the 2 person ball game. d/e) Sensor-level time-frequency spectra of a single OPM over the sensorimotor cortex of participant 2, during the touching task. Data are recorded without (d) and with (e) matrix coils active. Note how data, particularly in the 10-15 Hz band, are corrupted with artefact when the remnant field is not compensated. f) Comparison of task-induced oscillatory change across all frequencies, with (blue) and without (red) field nulling. Values represent percentage change from baseline. Again, results show that movement artefact masks alpha desynchronisation. Inset images show the location of the sensor.

To demonstrate the effectiveness of the coils, the two-person experiments, described above, were repeated without the coils activated. We expected that the strength and spatial variation of the magnetic field over the helmet would increase when the coils were switched off, such that the artefacts generated by movements would obfuscate the neural response.

Figure 3a shows the distribution of coil currents required to cancel the magnetic field experienced by the OPMs in the two helmets during the ball game task (the colour of each coil represents the amplitude of the applied current). Figures 3b and 3c show the level of field cancellation achieved over each helmet, for the guided touch and ball game tasks respectively. In both cases, the mean and standard deviation of the absolute values of the remnant magnetic fields, reported by the nulling sensors are shown. During the touch task the remnant field decreased from 3.3 ± 1.8 nT to 0.48 ± 0.44 nT (a factor of 6.9) for participant 1 and from 4.1 ± 2.2 nT to 0.43 ± 0.40 nT (a factor of 9.5) for participant 2. During the two-person ball game the field decreased from 3.8 ± 2.8 nT to 1.6 ± 1.3 nT (a factor of 2.4) for participant 1 and from 5.7 ± 2.9 nT to 2.0 ± 1.2 nT (a factor of 2.9) for participant 3.

To further demonstrate the effectiveness of the matrix coil system, we undertook a sensor level analysis. Figures 3d-e show trial averaged time frequency spectrograms from an OPM placed over the left motor cortex of participant 2, during the touching experiment (even trials). Panel d shows the case when the matrix coils were inactive and panel e shows equivalent data, from the same sensor, when the matrix coil system was used. with the system inactive, large positive changes from baseline spectral density extend across the alpha and beta bands. However, these become negative changes (reflecting the genuine task induced response) when the coil is activated. This degradation is caused by movements of the array through the non-zero remnant field. When activated, the matrix coil reduces the size of the artefacts, revealing the expected response. Figure 3f shows a comparison of the percentage change from baseline of measured oscillatory amplitude, in six key frequency bands, during the task (0.5 to 2 seconds). Mean values are shown, and error bars represent standard error over trials. Most strikingly, alpha desynchronisation is masked when the matrix coil system is not active. These effects show the need for magnetic field compensation during OPM-MEG experiments and highlight the performance and adaptability of the matrix coils.

### Solo demonstrations

To further demonstrate the effectiveness of the matrix coil system we conducted a series of ball-game experiments on a single participant (participant 2). 37 OPMs were spread over the helmet to obtain whole-head coverage. On receipt of an audio cue, the participant was instructed to bounce the table-tennis ball on the bat for 10 seconds, until a second audio cue instructed them to rest for 5 seconds. This process was repeated 40 times. The experiment was repeated with and without the matrix coil system activated. Movement of the helmet was recorded. To show the flexibility afforded by the matrix coil system, the participant conducted the task in three different positions: standing, seated on a chair at the centre of the coil, and seated on the floor.

The results show the expected decrease in beta oscillatory power in the left motor cortex during the periods when the participant was bouncing the table-tennis ball. Figures 4a-c show results for the standing, seated on a chair and seated on the floor condition, respectively. For each case, (i) shows the position of the participant during the task, (ii) shows beamformer images of beta modulation contrasting task (2 – 4 s) and rest (8 – 10 s), demonstrating activation of the motor cortices and (iii) shows the degree of DC magnetic field cancellation achieved at each position. (iv) shows the task induced (percentage) change from baseline of oscillatory amplitude in six key frequency bands in data from a single sensor sited over the left motor cortex. These indicate that the alpha band modulation was again obscured without field nulling. We also note the consistency of the field nulling and artefact reduction achieved at each of the three helmet locations.

**Figure 4:**
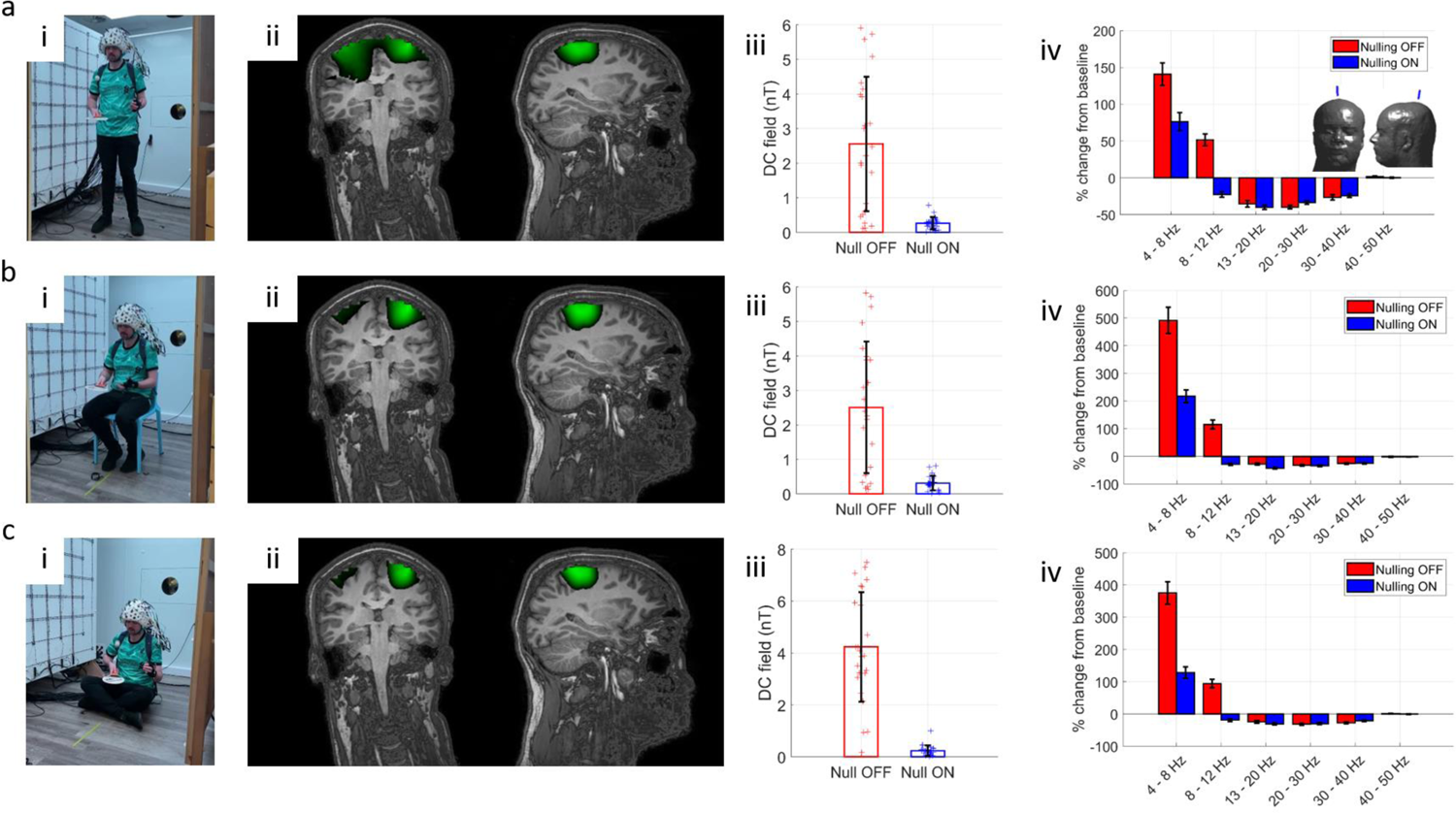
OPM-MEG data collected at various locations in the MSR, enabled by use of the matrix coils. The three rows show results for cases where the participant was stood up (a), sat on a chair (b) and sat on the floor (c), highlighting the wide range of scanning options afforded by matrix coils. i) Photos of a participant in each position. ii) Beamformer images showing the spatial signature of beta modulation (thresholded to 70% of the maximum value). The spatial pattern suggests activity in the sensorimotor regions when the participant bounces the ball on the bat. Note here all data are taken from the experiment where the matrix coils were active. iii) Bar charts showing the strength of the remnant DC magnetic field, with (blue) and without (red) nulling. iv) Task induced (percentage) change from baseline of neural oscillations in 6 key frequency bands (sensor-level results). As previously, movement artefact masks alpha desynchronisation. Inset shows the chosen sensor location. Photographs are of an author.

## 3. Discussion

Brain stimulation in functional imaging is often provided by artificial controlled events, which take place in restrictive, claustrophobic environments. Whilst useful, such experiments are of limited utility for understanding of how the human brain works in its native surroundings. Multi-modal stimuli, such as audio-visual footage and immersive (real or virtual) environments are now routinely deployed to investigate brain function during spontaneous, interactive events which more closely mimic real-life. Such naturalistic settings are crucial for collecting ecologically valid neuroscientific data; indeed it has been postulated that the evolution of the human brain is closely linked to a need for complex social interactions (Dunbar, 1998). For this reason, the importance of developing neuroimaging platforms that can interrogate brain function in naturalistic settings, is paramount. To date, the necessary technology to image brain function has been lacking either in performance or viability. Our work, for the first time, introduces a hyperscanning platform capable of direct detection of electrophysiological responses, with millisecond temporal and millimetre spatial precision, during natural, live interactions.

Our method was enabled by two technological advances, OPMs and matrix coils. Small and lightweight OPMs facilitate the high-precision measurement of magnetic fields generated by neuronal current. Unlike conventional MEG detectors, OPMs do not require cryogenic cooling and so can be placed closer to the head surface, improving data fidelity. Moreover, their lightweight nature allows sensors to move with the head. The flexibility of OPM-MEG has been clearly demonstrated here; an OPM array originally designed for a single participant was split across two helmets allowing simultaneous MEG data acquisition in two people. Accurate knowledge of sensor locations coupled with source analysis allows the derivation of functional images and interrogation of the time-frequency evolution of electrical activity. The synchronised nature of the recordings enables precise analysis of the relative timings of responses across participants. The second critical element of our system is the matrix coil system, which produces the zero-field environment in which the OPMs must be operated. The zero magnetic field requirement of OPMs is a significant barrier to producing wearable MEG systems which tolerate movement. OPMs have a low dynamic range and movement (even in an OPM-optimised MSR with a remnant field of a few nT) can render them inoperable. In addition, a changing field at the sensor induced by participant motion can generate marked changes in OPM sensor gain. Most importantly, movement generates large artefacts which obscure the neuromagnetic field. The use of the matrix coil ensured that OPMs remained operational and minimised gain changes in experiments. Further, motion induced artefacts were minimised; to an extent that, in the absence of matrix coil nulling, the expected alpha event related desynchronisation was completely masked by movement artefact – but recovered with the coil activated.

To demonstrate the adaptability of the matrix coil, we also performed single-person OPM-MEG recordings of a participant standing, seated on a chair and seated on the floor. The robustness of the spatial signature and temporal dynamics of the reconstructed neuronal activity highlights the utility of the system. It is significant that traditional Helmholtz or bi-planar coil designs are formed from fixed current paths which generate a known magnetic field within a prespecified volume. Such a design enables participant head movement within that volume but does not allow the nulled volume to be moved around the MSR. The matrix approach allows the coil to ‘re-design itself’, altering the current distribution in response to the location of the participant. This allows participants to be positioned anywhere between the coils, and even facilitates the creation of two separate nulled volumes, which is essential for hyperscanning. The modular nature of the matrix coil system also makes its design and construction simple compared to winding the intricate wire paths required by distributed coils systems: complexity is shifted to the coil amplifier and field control systems. Importantly, the data-driven field cancellation approach accounts for any helmet position, coil layout and the magnetic field distortions due to the presence of mu-metal, readily adapting to any size or shape of MSR.

There are some limitations of the present system to consider. Results showed that the performance of the matrix coil system during the two-person ball game was not as good as that during the guided touch or individual person paradigms. This is due to the increased separation of the two participants (required to hit the ball) which pushed the helmets worn by the participants to the edges of the coil planes. Consequently, fewer unit coils were available to contribute to the nulling and so performance was limited. During the standing experiments, the relative heights of the participants and their proximity to the upper-most section of coils had a similar effect. However, this limitation could be readily solved by expanding the coil array. In fact, extension of coil placement onto all six faces of the MSR would enable a wider variety of magnetic fields to be produced, increasing the range of possible experimental setups. Other extensions to our system include enabling the coil control software to account for low-frequency changes in the remnant magnetic field of the room (by imposing feedback controllers on the OPMs’ sensitive outputs, updating the coil calibration matrix either via optical tracking and calculation or by applying known oscillating currents to each coil). This would allow participant translations away from the initial nulled volume, paving the way for high-fidelity MEG acquisition in ambulatory participants.

The demonstrations presented in this paper show how hyperscanning can lead to novel findings. For example, our ball game paradigm reveals correlation of brain activity in two interacting participants. Nevertheless, these are simple demonstrations that only hint at the possibilities for OPM-MEG hyperscanning. Previous work has shown myriad possibilities: An excellent example is interactions between babies and their parents - indeed, past studies have employed EEG hyperscanning to show how the brains of a mother and baby demonstrate oscillatory synchronisation during normal social interactions, and that features of social interaction (e.g. eye contact) modulate the level of synchronisation (Leong et al., 2017). This prior work demonstrated the power of hyperscanning, but it was based upon technology that is limited (EEG is highly motion sensitive, spatial resolution is limited (particularly in infants where electrical potentials are distorted by the fontanelle) and high frequencies (beta and gamma oscillations) are disrupted by artefact). The OPM-MEG technology we have developed overcomes these limitations. Similarly, it offers possibilities for new clinical investigations, for example of social interaction in disorders such as autism.

Ultimately, a true understanding of the brain, and the many disorders that affect it, will only come through the ability to assess naturalistic function. Social interaction is a cornerstone of human development, and so understanding brain function during naturalistic interaction is a critical step along this path. OPM-MEG technology offers a means to do this, with an adaptable scanner equally able to scan one person or two people, using the same instrumentation, providing high fidelity measurements of brain activity. OPM-MEG using matrix coil technology thus has the potential to become the method of choice for future multi-person neuroimaging studies.

## 4. Methods

### OPM-MEG system

The OPM MEG system used here (excepting the matrix coils) is described in detail by Hill et al. (Hill et al., 2020); here we outline briefly its main features. The system (shown schematically in Figure 5) is housed inside a MSR which is optimised for OPM operation (MuRoom, Magnetic Shields Limited, Kent UK). The MSR features 4 layers of mu-metal and 1 layer of copper, along with demagnetisation coils (Altarev et al., 2015; Voigt et al., 2013). The typical remnant magnetic fields and field gradients at the centre of the room are of order 2 nT and 2 nT/m, respectively. Note however that for the experiments carried out here, the head was not positioned at the centre of the room and the remnant fields are consequently larger, as can be seen in Figure 4.

**Figure 5:**
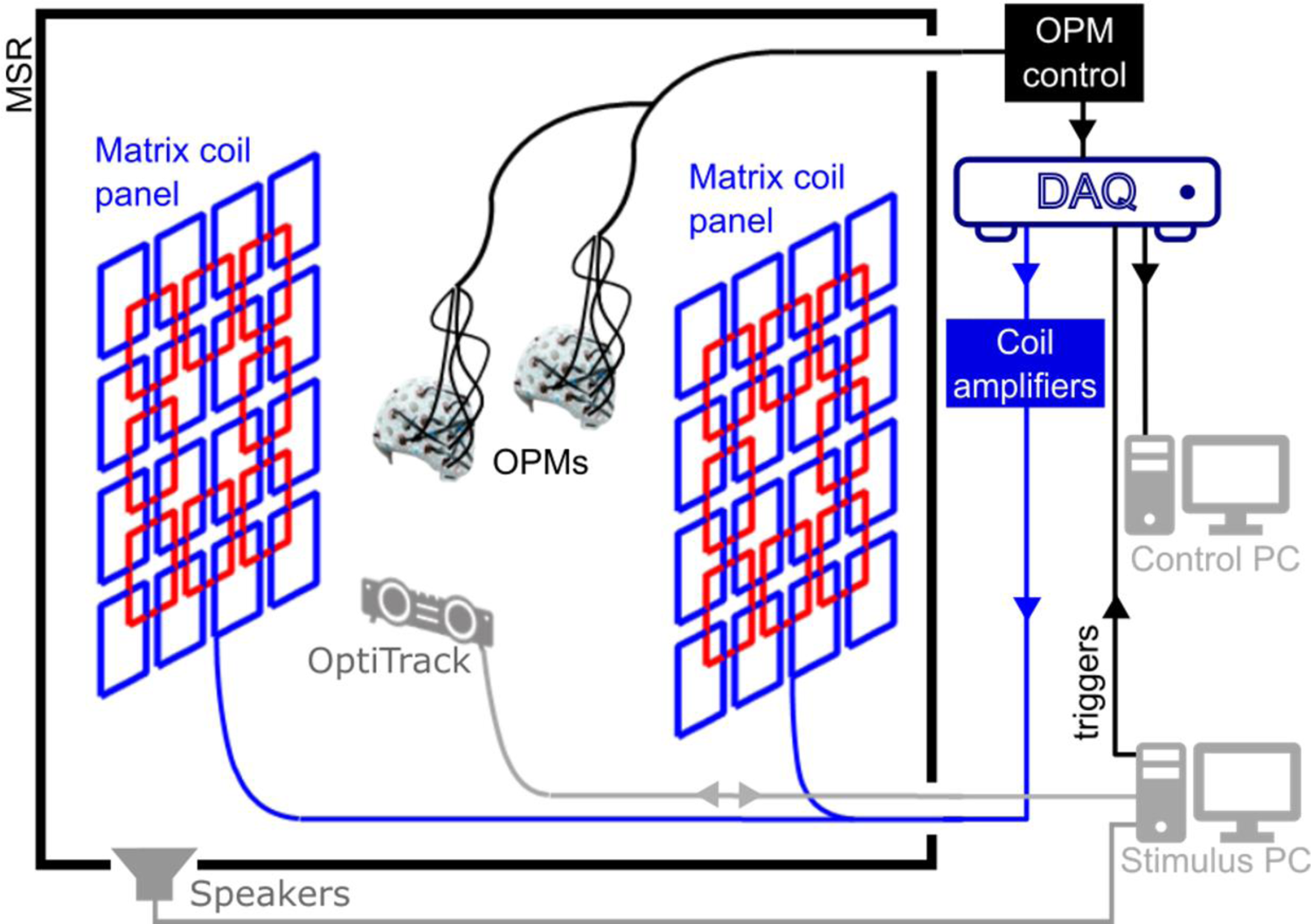
OPM-MEG system schematic. The system is housed in a magnetically shielded room (MSR). OPMs are interfaced with a series of data acquisition devices. Data from the OPMs are used to drive the matrix coil field nulling process, before a MEG recording begins. Optical tracking of the helmets is performed to monitor motion during a session. Instruction is passed to the participants via auditory cues controlled using a separate stimulus PC.

Up to 50, second generation, QuSpin Inc. (Colorado, USA) zero-field magnetometers are available for array formation (see Tierney et al. (Tierney et al., 2019) for a review of OPM physics and Osborne et al. (Shah et al., 2018) for specific details of the QuSpin sensor). The OPMs were mounted inside 3D-printed, rigid scanner-casts which allow co-registration of OPM positions and orientations to anatomical MRI’s (whole-head MRI scans were generated using a 3 T Philips Ingenia system, running an MPRAGE sequence, at an isotropic spatial resolution of 1 mm) of the participants’ heads (Boto et al., 2017; Hill et al., 2020; Zetter et al., 2019). OPMs were configured to record the component of magnetic field which is radial to the surface of the head. OPM data were sampled at 1,200 Hz using a series of National Instruments (NI, Texas, USA) NI-9205 16-bit analogue to digital converters interfaced with LabVIEW (NI, Texas, USA). Since all the OPMs are sampled and controlled using the same equipment, no additional timing signals or hardware are required to synchronise the data collected from the two helmets. Participant movements were tracked using a OptiTrack V120:Duo (NaturalPoint Inc., Corvalis, USA) optical tracking system which provides sub-1-millimetre and sub-1-degree precision optical tracking of multiple rigid bodies at a sample rate of 120 Hz. Two cameras, each with an array of 15 infrared (IR) LEDs, are used to illuminate IR reflective markers and the combined coordinates of multiple markers are used to form a rigid body tracking with 6 degrees of freedom (x, y and z translations, pitch, yaw and roll rotations).

### Matrix coils

Our aim was to develop a system that produces a magnetic field, equal in magnitude but opposite in direction to the remnant magnetic field within target volume(s) inside the MSR, thereby nulling the field. Matrix-coil systems feature an array of small, simple, unit coils positioned around the participant. Superposition of the magnetic field generated by multiple coils, each carrying an independently controllable current, enables the production of arbitrary patterns of magnetic field variation within a selected target volume (Garda and Galias, 2014; Juchem et al., 2010). Similar multi-coil shimming systems have been developed for MRI (Juchem et al., 2015, 2011). Our matrix coil system was constructed using a bi-planar design, with each plane containing 24 square coils (square side length 38 cm). The coils are arranged on a 4 x 4 grid with an overlapping 3 x 3 grid in which the central coil is omitted (Figure 3a). Each coil was wound by hand using 10 turns of 0.56 mm diameter copper wire, tightly wrapped around a series of plastic guides attached to a wooden structure (coil resistance ∼2 Ω, coil inductance ∼160 µH). The two planes are sited on either side of the participant(s), separated by 150 cm.

Each unit coil is connected to a single output of a 48-channel, low-noise, voltage amplifier that was designed and constructed in-house. This is interfaced to three NI-9264 16-bit, digital to analogue converters (DACs) that are controlled using LabVIEW. The voltages applied at the amplifier input range between ±10 V (least significant bit (lsb) voltage = 20 V/2^16^ = 0.305 mV). The maximum electrical current in the coil is tuned by an additional series resistance, which in this setup was 1.2 kΩ, chosen such that the magnetic field noise generated by the coils was beneath the noise floor of the OPMs. The coil driver current noise at this resistance is <10 nA/√Hz in the 1-100 Hz band, we estimate this translates to <20 fT/√Hz noise in the field from all 48 coils at the centre of the planes (see supplementary material), for comparison, the OPM noise floor is <10 fT/√Hz in this frequency range so the two are comparable. The maximum current which can be applied to each coil is ±8.33 mA, and the lsb current is 2.54 µA.

To null the remnant magnetic field inside the MSR during a MEG experiment we employed a data-driven approach. If the magnetic field measured by the *n*^th^ OPM in an array of *N* sensors due to unit current in the *m*^th^ coil in a set of *M* (= 48) matrix coils is written as db^n^⁄dI_m_’, we can form a (*N* x *M*) coil calibration matrix, *A*, from the full set of values. The field nulling problem can then be described using the following matrix equation:

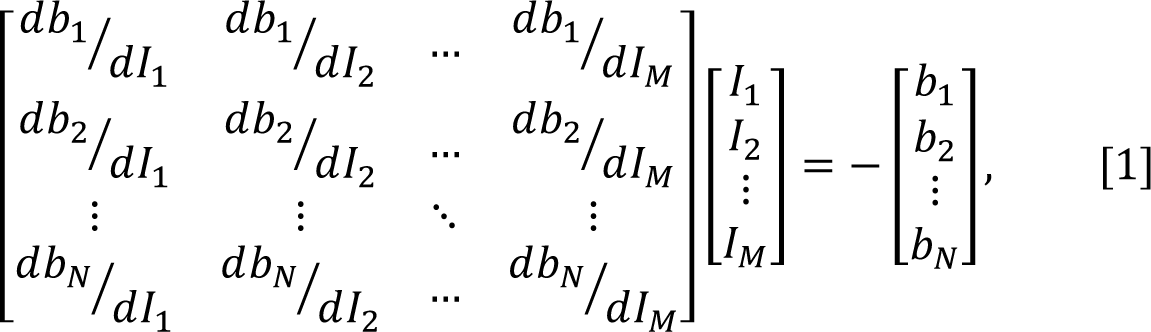

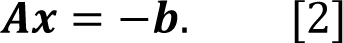

where the (*M* x 1) column vector *x* contains the currents applied to each coil and the (*N* x 1) column vector *b* characterises the magnetic field to be cancelled. *b* is formed using the DC field values measured at the sensors, the negative sign is used to ensure the calculated currents null the magnetic field measured by the array.

The coil currents required to minimise the sum of squares of the measured magnetic field values can be found by identifying the negative of the Moore-Penrose pseudo-inverse matrix of *A*,

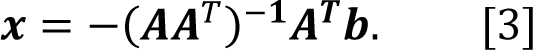

To minimise the power dissipated by the system, and ensure the solution is physically realisable, the matrix *AA*^T^ can be regularised prior to inversion by addition of a matrix αI where I is the identity matrix of the same dimensions as *AA*^T^ and α is a regularisation parameter i.e.

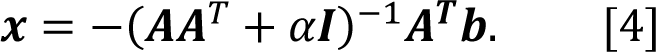

To keep the coil currents within the allowed bounds, equation [4] is cast as a feed-forward controller: coil currents are incrementally updated, based on the OPM field measurements at each timepoint *i*, and the currents applied at the preceding time point as

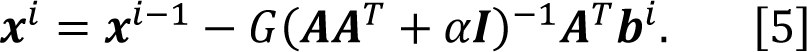

The gain coefficient G is empirically set to produce a stable reduction of the measured fields towards zero on a timescale of a few seconds.

This approach can readily be adapted to changes in the number and shape of the unit coils and flexibly incorporates multiple sensor arrays. However, it only considers the field values at the sensor positions and as a result, unwanted deviations in magnetic field could occur between target points. Coil calibration data for populating the matrix, *A*, can be collected in a variety of ways depending on the available sensing technology e.g. by pulsing each coil in turn or by applying a known sinusoidal current to each coil. Values could also be calculated based on known sensor positions, coil design and geometry of the MSR.

The nulling procedure described above was implemented in LabVIEW. Participants were instructed to remain still whilst a 5 V (4.16 mA), 100 ms pulse was applied to each coil in turn. The change in field experienced by each OPM was measured by interfacing the OPMs with LabVIEW and operating the sensors in their field-zeroing mode. In this mode, QuSpin OPMs can measure the DC magnetic field experienced by the cell (along two orthogonal directions) with a dynamic range of ±50 nT (Shah et al., 2018; Shah and Hughes, 2015). The field zeroing procedure is generally carried out prior to OPM gain calibration when an experiment is performed, providing measurements of the offset magnetic fields required to produce the zero-field environment in the sensor cell. The regularisation parameter α (in Eq. [5]) was set to 1% of the maximum singular value of the matrix *AA*^T^. The feed-forward controller gain was set to 0.1 with a time step of 100 ms.

The time needed for the calibration process scales with the number of coils and takes around 1 minute to complete for the 48-coil system. The final coil currents were held constant during the experiments, i.e. no dynamic tracking of changes in magnetic field was applied (we note the magnetic field drift in our MSR is on the order of 200 pT over 10 minutes). The LabVIEW program stores the magnetic field values reported by each sensor prior to calibration, along with the coil calibration matrix, the final voltages applied to each coil and the final magnetic field values.

### Data acquisition

All data were collected by the authors. Participants provided written informed consent for all experiments. All studies were approved by the University of Nottingham’s Faculty of Medicine and Health Sciences Research Ethics Committee. Additional guidelines to mitigate the risk of transmission of COVID-19 were adhered to by all participants and experimenters: participants wore face masks and visors during the two-person experiments as can be seen in Figures 1 and 2. Audio cues to structure the experiment were single beeps, generated by MATLAB (MathWorks, CA, USA), and played through speakers placed inside waveguides in the top corners of the MSR. MATLAB was also used to generate a trigger signal at the same time as the audio cues which was recorded along with the OPM data for synchronisation.

### Guided touch task

For the two-person touching task, each participant wore an array of 16 OPMs mounted in a 3D-printed scanner-cast. Participant 1 (female, aged 30, height 172 cm) wore a scanner-cast which was custom-made for their head based on an anatomical MRI (Chalk Studios, London, UK) meaning that co-registration of the positions and orientations of the OPMs with respect to the participant’s brain was known(Boto et al., 2017). Participant 2 (male, aged 25, height 182 cm) wore a rigid, additively manufactured generic scanner-cast (Added Scientific Limited, Nottingham, UK) which was designed to fit an average adult head-shape (Hill et al., 2020). The co-registration of OPM sensors to the anatomy of participant 2 was performed by using 3D structured light scans combined with the known structure of the generic helmet (Hill et al., 2020; Zetter et al., 2019). The positions and orientations of the OPMs used in the experiments for each participant are shown in Figure 6, with the sensors used for field nulling highlighted. Sensors were arranged to cover the left hemisphere with additional sensors placed at the front, back and right sides of the head to extend the region of space over which fields were considered in the nulling process. During the experiment, each participant was instructed to reach over and stroke the right hand of the other participant with their right hand, following an audio cue. The audio cue repeated every 5 seconds and the active participant alternated between trials. Each participant conducted 30 active trials.

**Figure 6:**
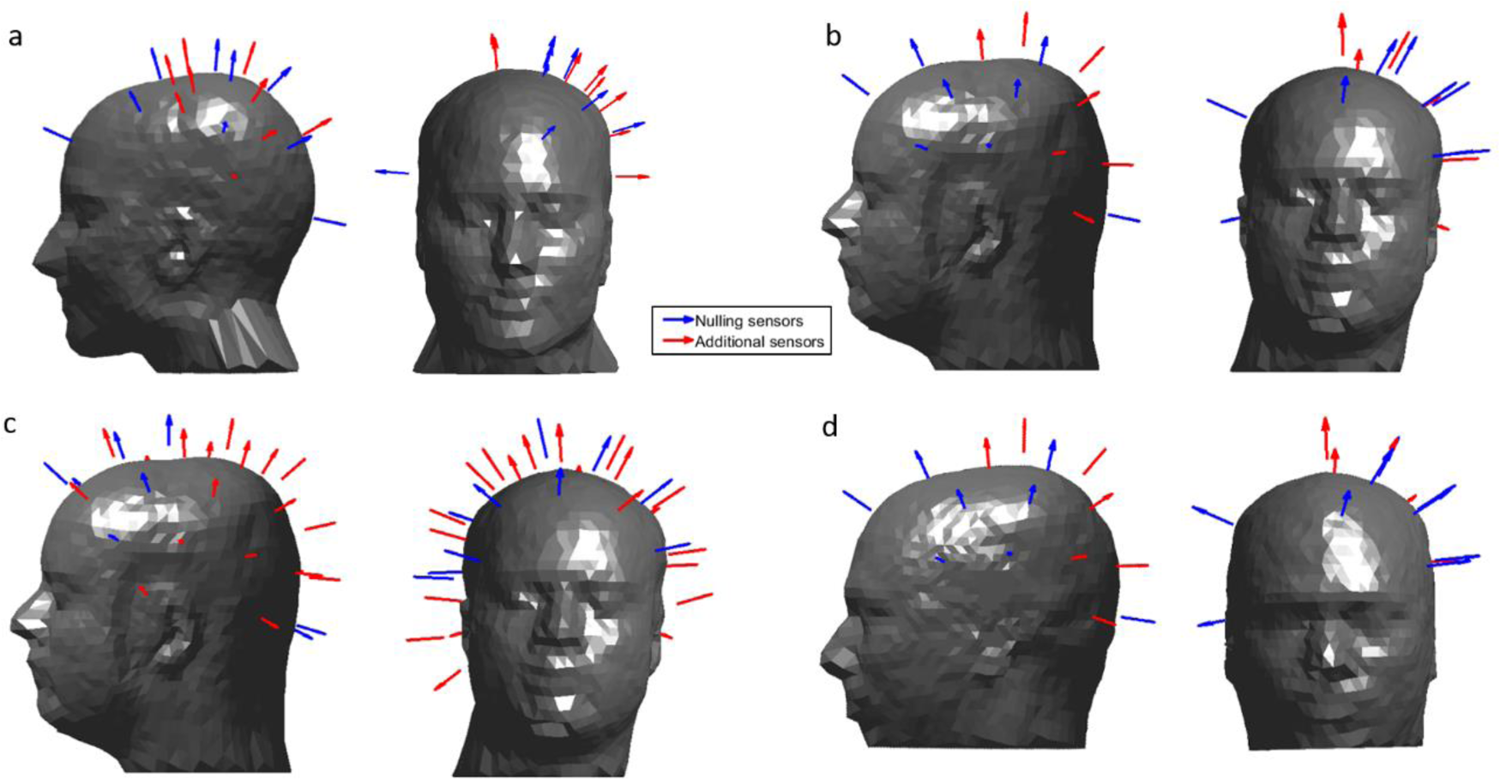
The position and orientation of the OPMs used in each experiment. Sensors which were used to inform the field nulling are shown in blue and additional sensors are shown in red. During the hyperscanning experiments, sensors were concentrated over the left sensorimotor cortex, with additional sensors placed at the right and the front of the head, to inform the nulling process. Coverage was extended over both hemispheres for the solo ball game. a) Sensor layout for participant 1 during the two-person touch and ball game tasks. b) Sensor layout for participant 2 during the two-person touching task. c) Sensor layout for participant 2 during the solo ball-game task (note coverage of both hemispheres). d) Sensor layout for participant 3 during the two-person ball game.

### Ball game task

During the two-person ball game experiment, participant 1 again wore an individualised scanner cast, whilst participant 3 (and male, aged 41, height 188 cm) wore a generic 3D printed helmet (co-registration as above). The participants were instructed to hit a table-tennis ball back and forth to each other for 5 seconds, following an audio cue. A second audio cue instructed the participants to stop their rally and rest for 7 seconds. This was repeated 25 times. Movement of the two helmets was again tracked using the OptiTrack camera system throughout the experiment. Each Hyperscanning task was repeated twice, with and without the matrix coils active (the participants were not blinded to this condition). Trials where the ball was dropped were noted and excluded from data analysis (two dropped balls in each condition).

### Solo experiments

For the solo MEG experiments, a single participant (participant 2) wore the generic scanner-cast containing 37 OPMs distributed over the entire head. During the experiment, the participant was instructed to bounce a table-tennis ball on the bat for 10 seconds following an audio cue. A second audio cue instructed the participants to stop and rest for 5 seconds. This was repeated 40 times.

Recordings were made while the participant stood up at the centre of the coil planes, then sat on a chair and finally sat on the floor of the MSR. The entire process was repeated with and without the matrix coils active. All trials were complete successfully without dropping a ball.

### Field nulling

In the hyperscanning experiments, measurements of the amplitude of two field components from 10 OPMs operating in field zeroing mode housed in the scanner-casts of each participant (i.e. the matrix *A* contains values from 20 OPMs giving *N* = 40 measurements in total) were used as inputs to the LabVIEW-based field nulling program described above. The 10 sensors on each helmet that were used for the nulling process, included the additional sensors sited at the front, back and right-hand sides of the head, as well as seven sensors sited over the left side of the head. Participants were asked to remain still whilst the system was calibrated and instructed to keep their feet planted throughout the experiment to avoid translating their heads away from the nulled volume.

During the solo experiments, The two field components measured by 12 OPMs (*N* = 24 total measurements) operating in field zeroing mode were used as inputs to the field nulling program. Field nulling sensors were chosen such that they spanned the full volume of the head and are highlighted in Figure 6.

### OPM settings

All OPM data were collected at a sample rate of 1,200 Hz using the equipment described earlier. Once the matrix coil currents had been set, the OPMs were field zeroed and calibrated using the QuSpin software. The OPMs were then set to their 0.33x gain mode (voltage to magnetic field conversion factor 0.9 V/nT) in which their dynamic range is ±5 nT. The default gain setting has a lower dynamic range of ±1.5 nT (2.7 V/nT) which was not used here as the outputs would have quickly saturated when participants moved during the nulling-off experiments.

### Data analysis

All code for analysis was custom written by the authors using MATLAB.

### Source reconstruction

A beamformer approach was used to generate the images shown in Figures 1, 2 and 5. An estimate of the neuronal current dipole strength, Q^_θ_(t), at time t and a position and orientation θ in the brain is formed via a weighted sum of the measured data as

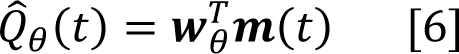

where *m*(t) is a vector containing the magnetic field measurements recorded by all OPMs and W_θ_ is a weights vector tuned to θ. The weights are chosen such that

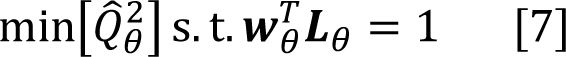

where L_θ_ is the forward field vector containing the solutions to the MEG forward problem for a unit dipole at θ. The optimal weights vector is expressed as

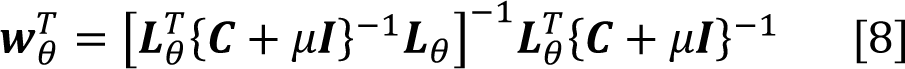

where C is the sensor data covariance matrix. Inversion of the covariance matrix is aided by Tikhonov regularisation (i.e. by addition of the identity matrix scaled by regularisation parameter μ).

To compute the weights vectors for each experiment, the entire dataset was filtered to the beta band (13 – 30 Hz) and used to compute the covariance matrix. The regularisation parameter μ was set to 0.01 times the leading singular value of the covariance matrix. The forward field vector was calculated using a multi-sphere head model and the current dipole approximation (Sarvas, 1987).

Images of activation show the pseudo-T-statistical contrast between data recorded in active and control windows. Specifically, two covariance matrices were computed for the active and control periods, C_a_ and C_c_ respectively, and the pseudo-T-statistical contrast, at θ, calculated as

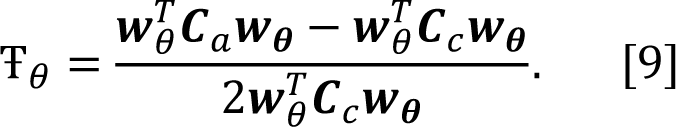

Pseudo-T-statistics were computed at the vertices of a regular 4 mm grid spanning the whole brain. This grid of Ŧ values was then thresholded to a percentage of the maximum value and overlaid onto the anatomical MRI of each participant. In the touch experiment the Ŧ values were thresholded to 80% of the maximum value for each condition. The active period was 0.5 to 2 seconds and the control period was 3 to 4.5 seconds. In the two-person ball game the active period was 2 to 4 seconds and the control period was 10 to 12 seconds. In the single-person ball game the Ŧ values were thresholded to 70% of the maximum value, the active period was 2 to 4 seconds and the control period was 8 to 10 seconds.

For each experiment, the location of the voxel with largest pseudo-T-statistic was determined for each participant. The signal from this peak location was reconstructed to form a ‘virtual electrode’ timecourse (using Equation 6), with beamformer weights calculated in the broad-band (1‒150 Hz).

The time-frequency spectra were generated by filtering this timecourse sequentially into overlapping frequency bands. For each band, the Hilbert envelope was calculated before segmenting and averaging over trials and concatenating in the frequency domain. The mean envelope in the beta band was computed using the virtual electrode timecourse filtered to the beta band (13‒30 Hz).

### Sensor-level analysis

Similar analysis was performed to generate the sensor level time-frequency spectra shown in Figure 3d and 3e. The bar charts comparing change from baseline activity in Figure 3f and 4abc (iv) were computed using contrasting active and control periods, as above, and non-overlapping frequency bands.

### Correlating activity

Correlation of brain activity (Figure 2e) was computed using the 3-6.5 s window of the beta band filtered virtual electrode average trial timecourse. The normalised, unbiased, autocorrelation and cross-correlation of the two timecourses were computed for a maximum lag of 3.5 s (i.e. the full duration of the data segment).

## Acknowledgements

We express our sincere thanks to collaborators at the Wellcome Centre for Human Neuroimaging, University College London, UK for extremely helpful discussions and ongoing support. We also thank Magnetic Shields Limited who designed and constructed our OPM-optimised magnetically shielded room. This work was supported by the UK Quantum Technology Hub in Sensing and Timing, funded by the Engineering and Physical Sciences Research Council (EPSRC) (EP/T001046/1), a Wellcome Collaborative Award in Science (203257/Z/16/Z and 203257/B/16/Z) and National Institutes of Health grant R01EB028772.

## Data and code availability statement

The raw sensor-level data collected in this work (without associated MRIs) will be made publicly available. The authors declare the following competing interests: V.S. is the founding director of QuSpin Inc., the commercial entity selling OPM magnetometers. J.O. is an employee of QuSpin. E.B. and M.J.B. are directors of Cerca Magnetics Limited, a spin-out company whose aim is to commercialise aspects of OPM-MEG technology. E.B., M.J.B., R.B., N.H. and R.H. hold founding equity in Cerca Magnetics Limited. N.H, P.G, M.J.B., and R.B. declare that they have a patent pending to the UK Government Intellectual Property Office (Application No. GB2109459.4) regarding the active magnetic shielding systems described in this work.

## Supplementary material

### 1. Motion during experiments

To assess the range of movements during the experiments and their comparability we further analysed the data recorded by the OptiTrack camera system. The helmet translation (x, y and z, defined relative to the camera position) and rotation (pitch roll and yaw, defined as rotation about the centre of mass of the rigid body which is formed from four infrared reflective markers placed onto the front or back of the helmet worn by the participant, depending on the field-of-view of the camera) data were segmented into trials. The first data point was subtracted from each trial such that the motion data reflected the *change* in translation or rotation of the rigid body on the helmet relative to its initial position during a trial. All six motion parameters are plotted in Figures S1-S7 for each experiment, for each trial and with and without field nulling to show the range of motions required in each task and comparability of these motions across conditions.

**Figure S1:**
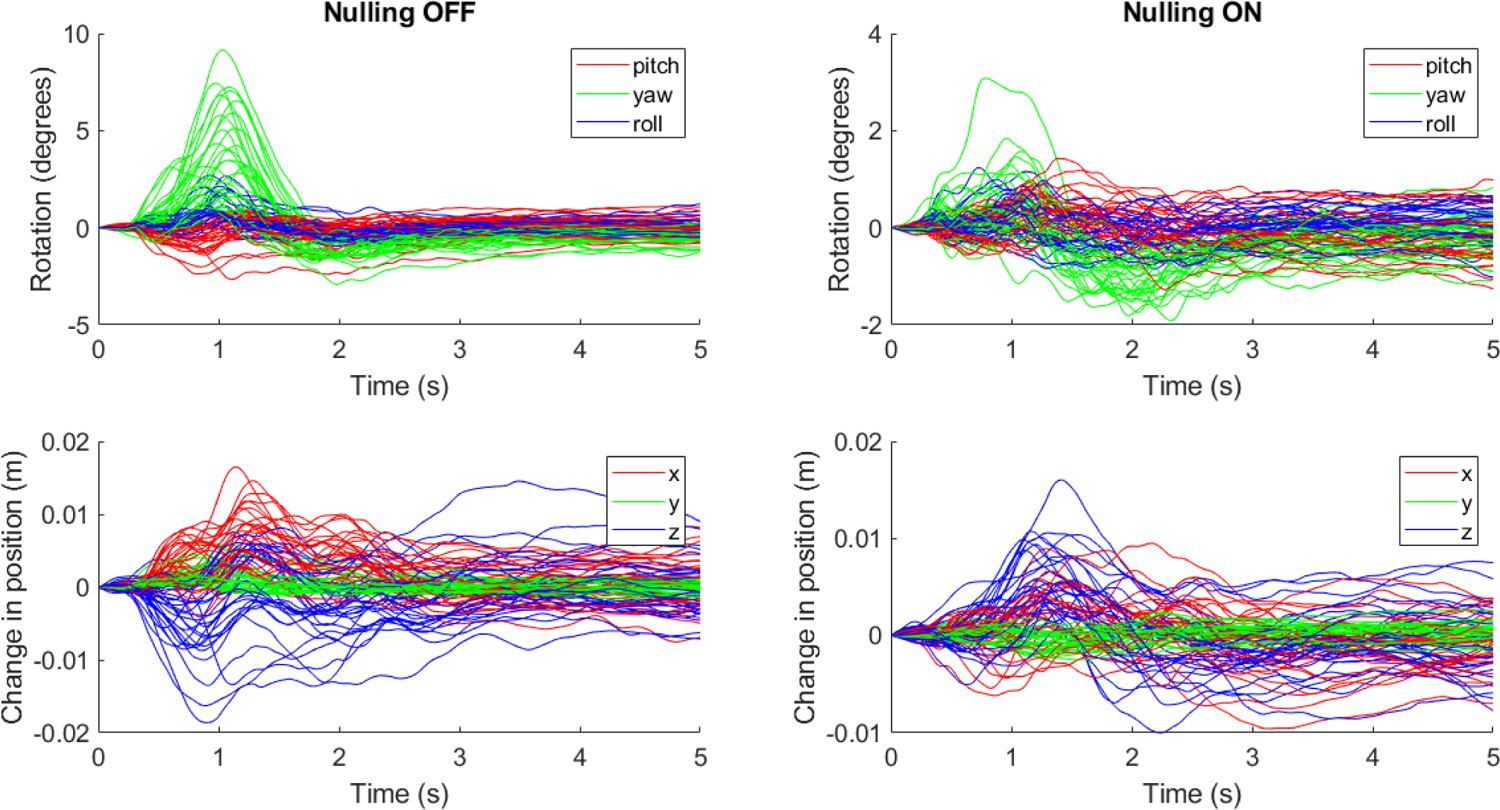
Change in the position and rotation of the helmet worn by participant 1 during the hyperscanning guided-touch experiment for trials in which participant 1 was active (i.e. leaning over the table and touching, as opposed to remaining still and being touched) repeated with and without the matrix coils activated. Top row shows change in helmet rotation about the centre of mass of the rigid body formed of 4 infrared reflective markers attached to the OPM-MEG helmet. Bottom row shows change in position of the centre of mass of the rigid body.

**Figure S2:**
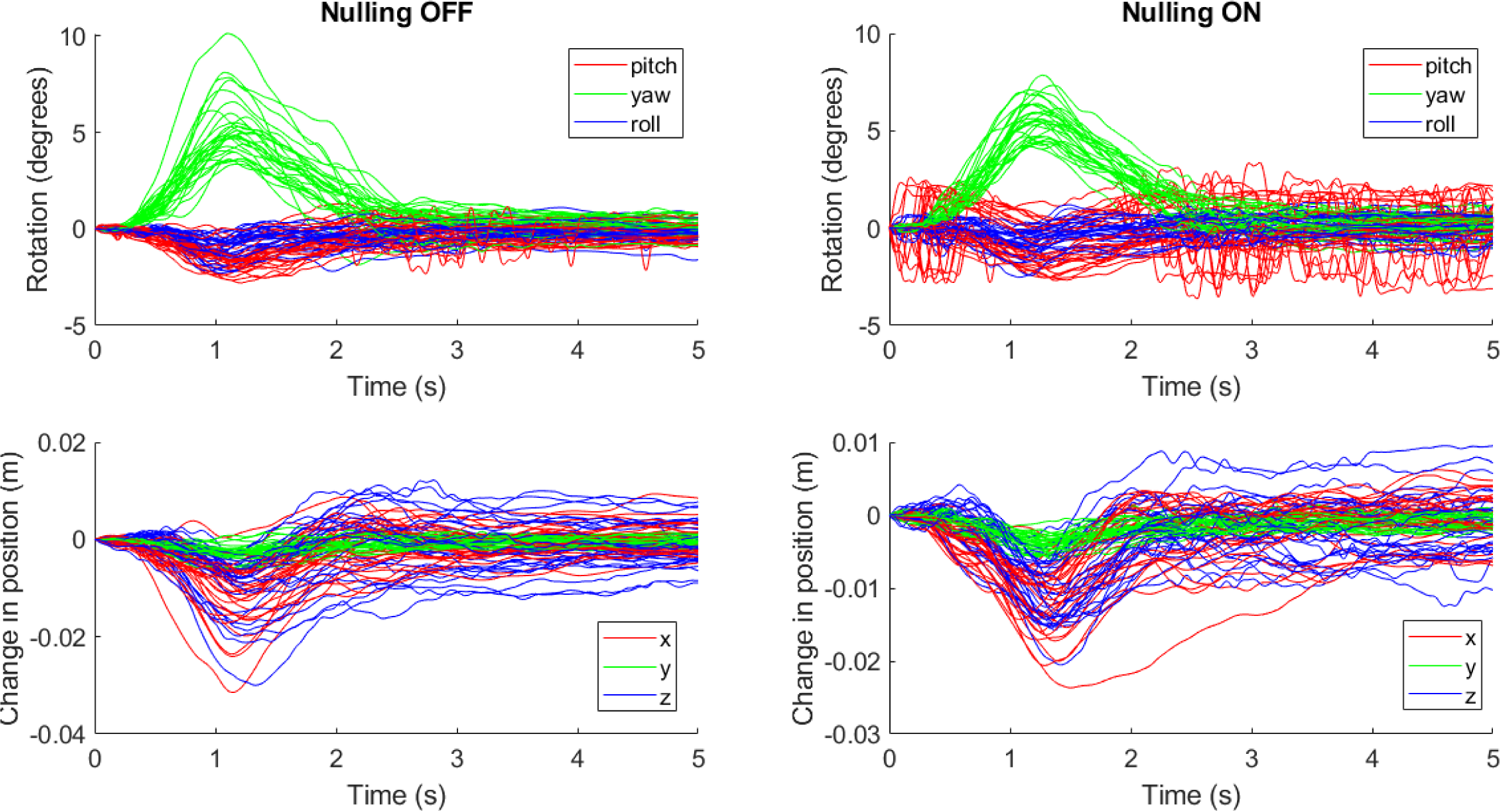
Change in the position and rotation of the helmet worn by participant 2 during the hyperscanning guided-touch experiment for trials in which participant 2 was active repeated with and without matrix coils activated.

**Figure S3:**
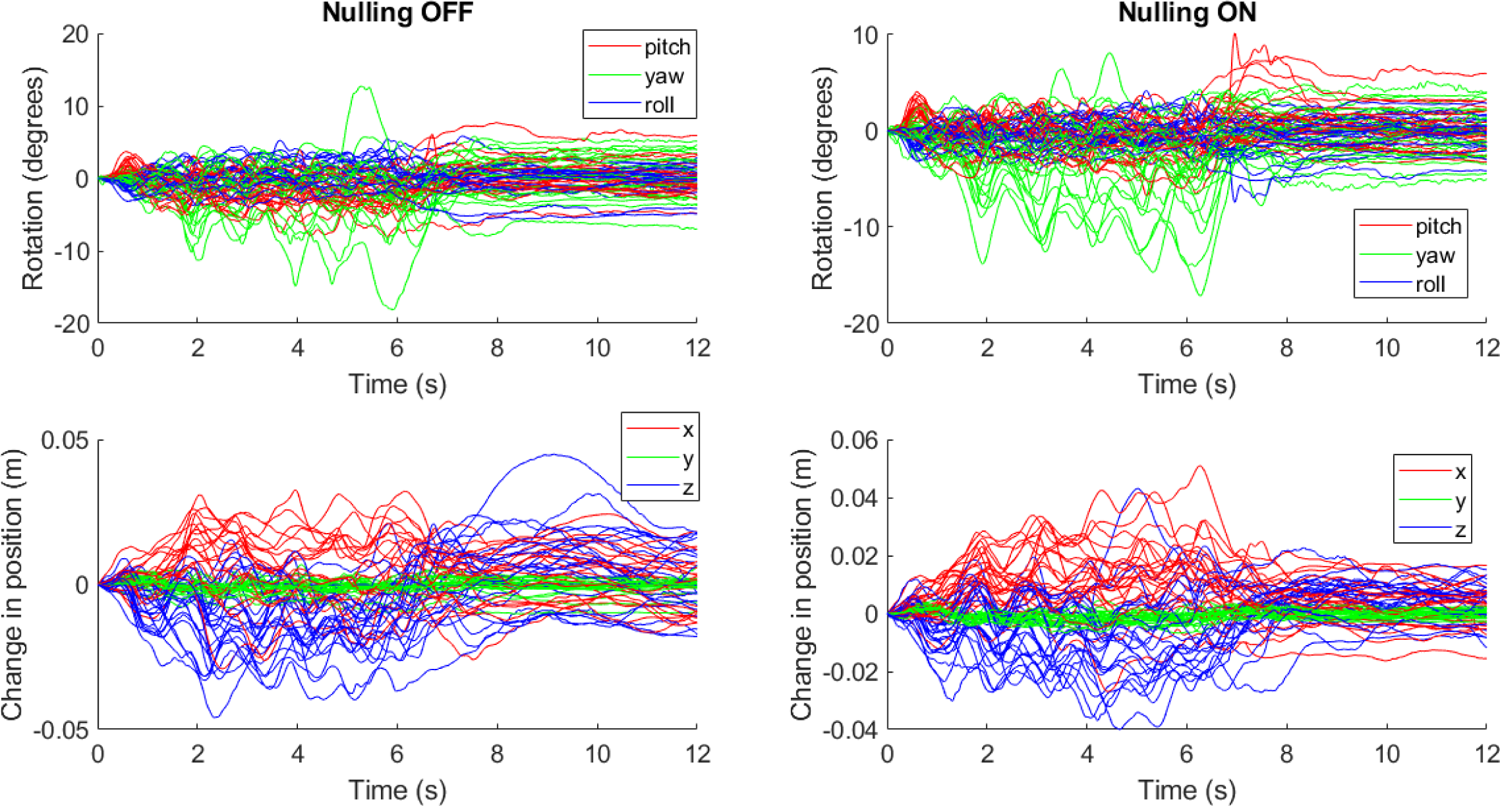
Change in the position and rotation of the helmet worn by participant 1 during the hyperscanning ball game experiment repeated with and without matrix coils activated.

**Figure S4:**
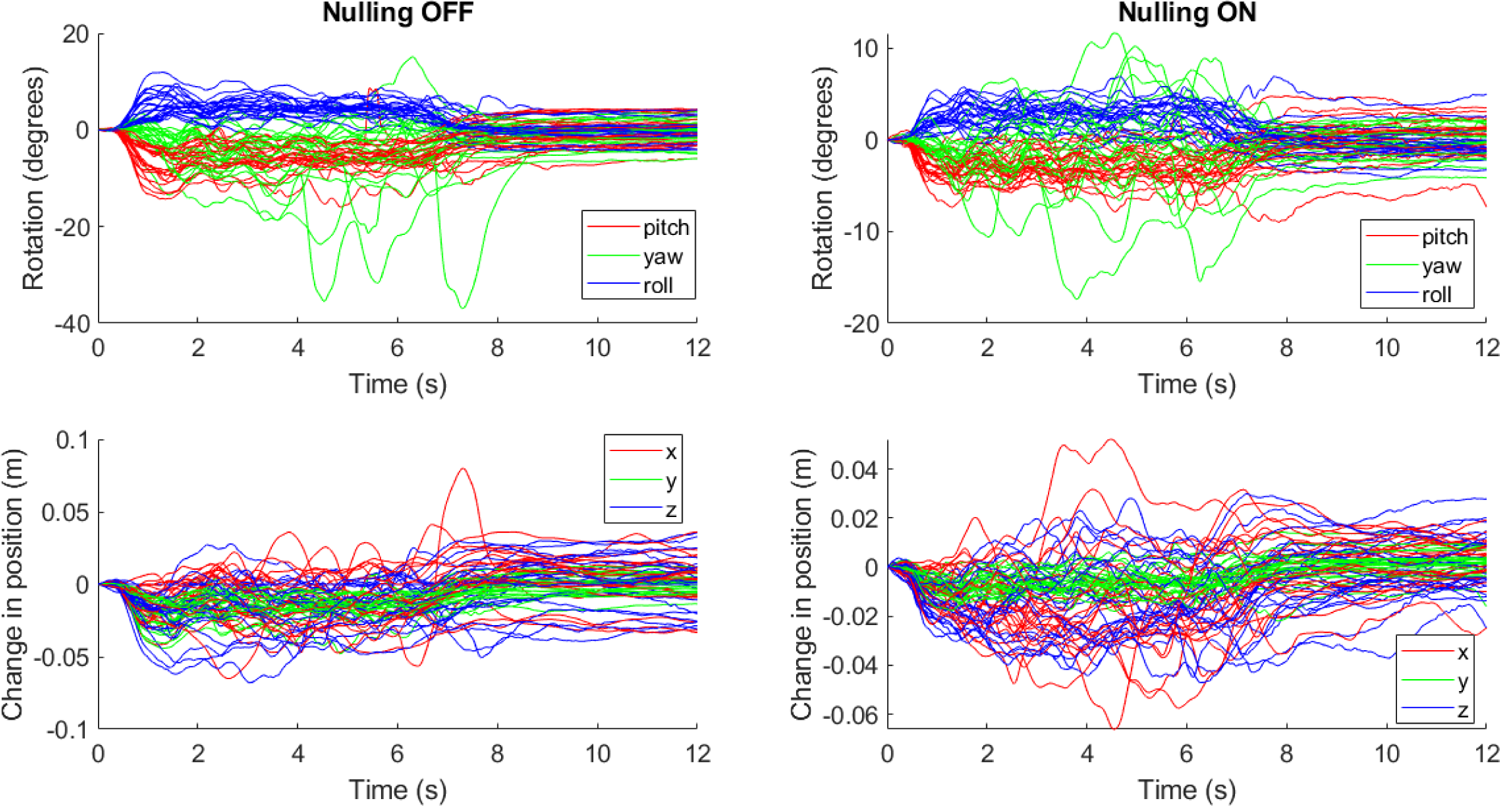
Change in the position and rotation of the helmet worn by participant 3 during the hyperscanning ball game experiment repeated with and without matrix coils activated.

**Figure S5:**
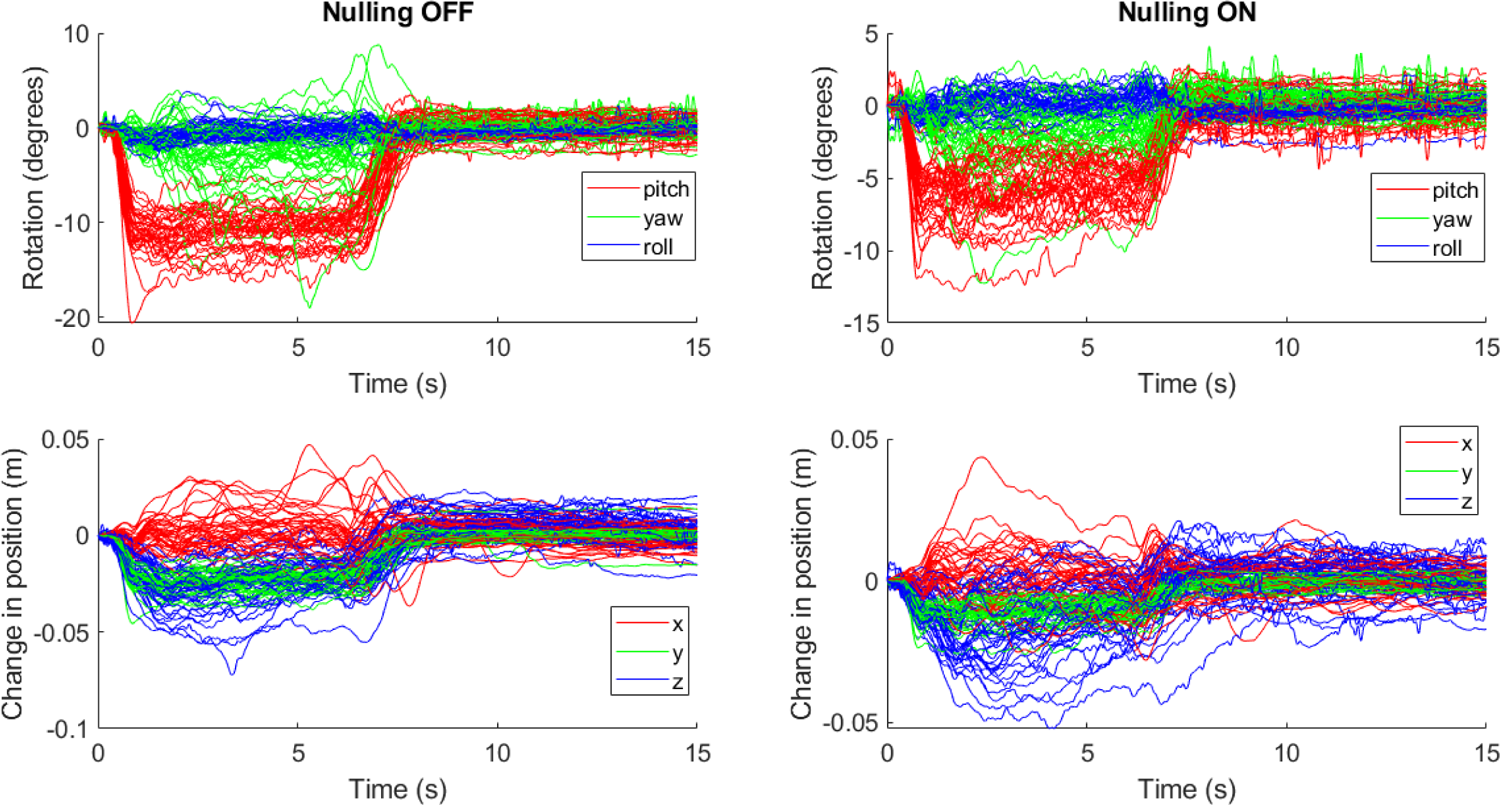
Change in the position and rotation of the helmet worn by participant 2 whilst stood up during the solo ball game experiment repeated with and without matrix coils activated.

**Figure S6:**
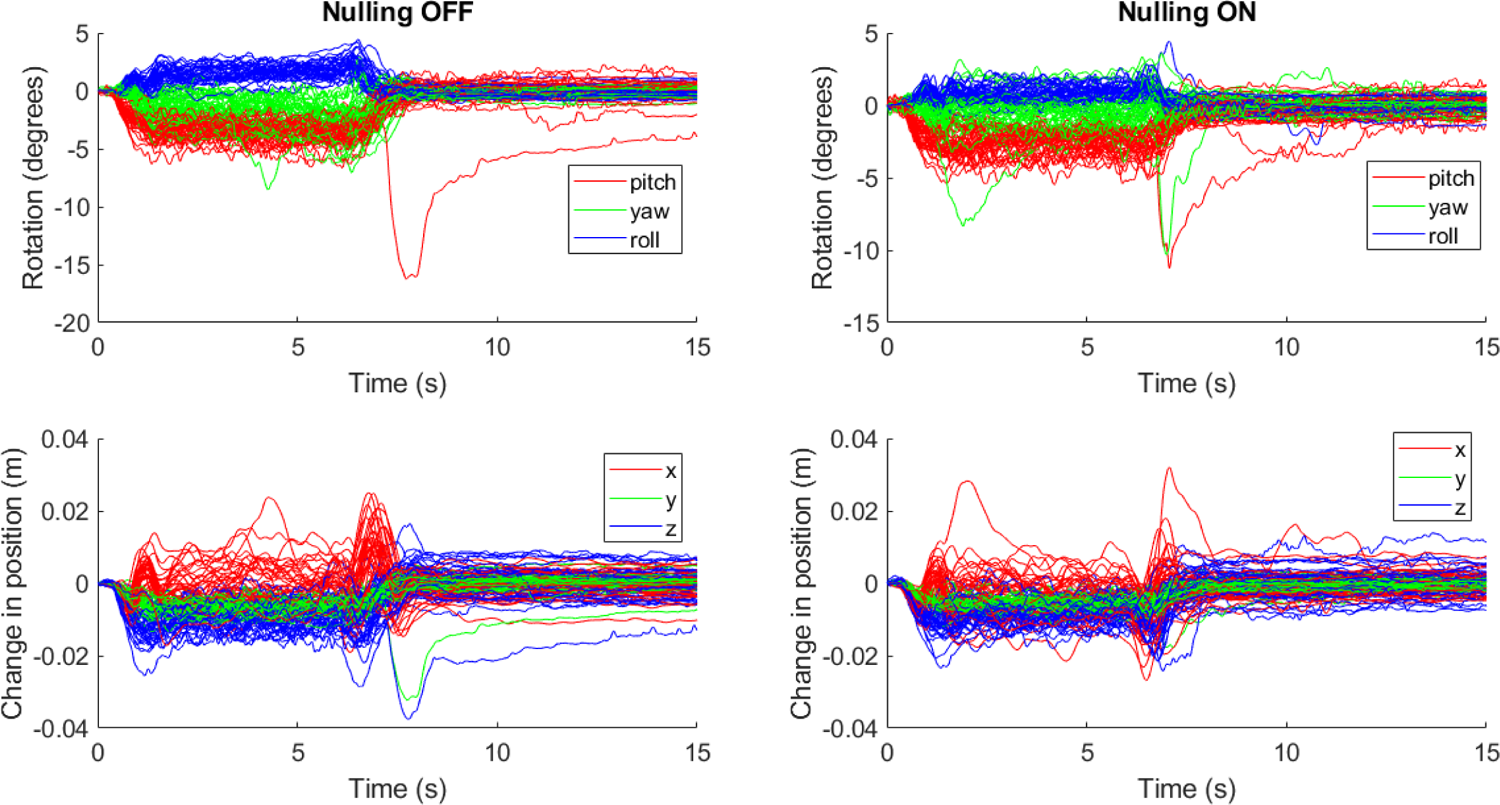
Change in the position and rotation of the helmet worn by participant 2 when seated on the chair during the solo ball game experiment repeated with and without matrix coils activated.

**Figure S7:**
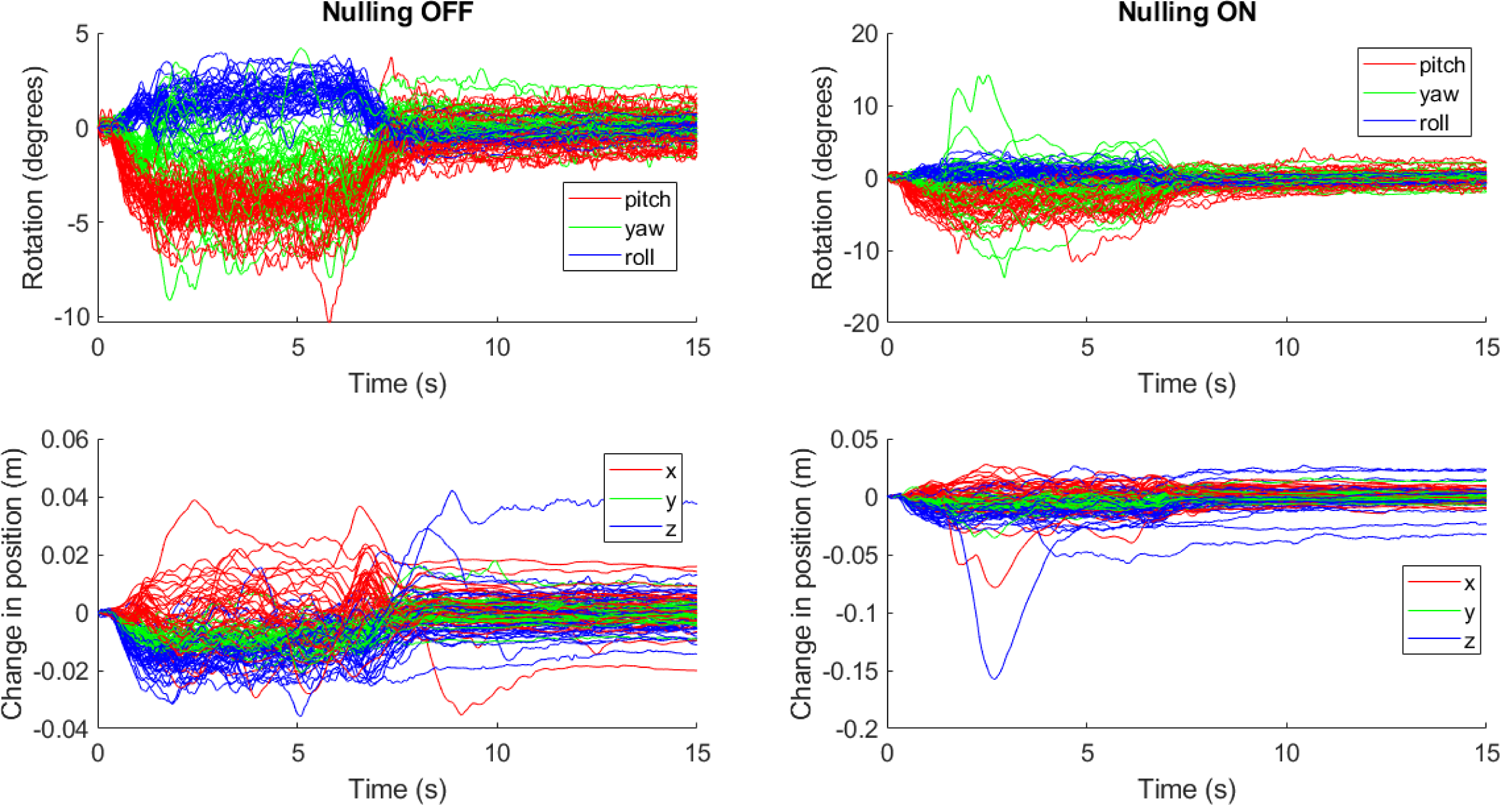
Change in the position and rotation of the helmet worn by participant 2 when seated on the floor during the solo ball game experiment repeated with and without matrix coils activated.

### 2. Matrix coil noise estimates

As the magnetic field generated by each of the 48 coils in the matrix coil will sum, careful consideration of the magnetic field noise induced by current noise in the system is required. A balance is needed between the maximum current which can be produced in order to generate the required magnetic field profiles, and field noise induced by the amplifier electronics that could mask brain activity.

The voltage across the 1.2 kΩ resistor in series with each coil with the voltage amplifiers active was recorded and converted to current. The current noise measured was <10 nA/√Hz in the 1-100 Hz band. We simulated the magnetic field (taking into account interactions between the magnetic fields generated by the coils and the high-permeability material used to construct the MSR) for 10 nA of current in each coil at each point in a regular 10-cm resolution grid spanning a 1.4 x 1.4 x 1.4 m^3^ volume between the matrix coil planes as shown in Figure S8. For each vector component of the magnetic field at each point in the grid we computed the square root of the sum of the squared magnetic field values for each coil as an estimate of the noise in each direction. The square root of the sum of these squared vector values was then taken as an estimate of the magnitude of the noise. Each noise estimate is expressed in units of fT/√Hz.

**Figure S8:**
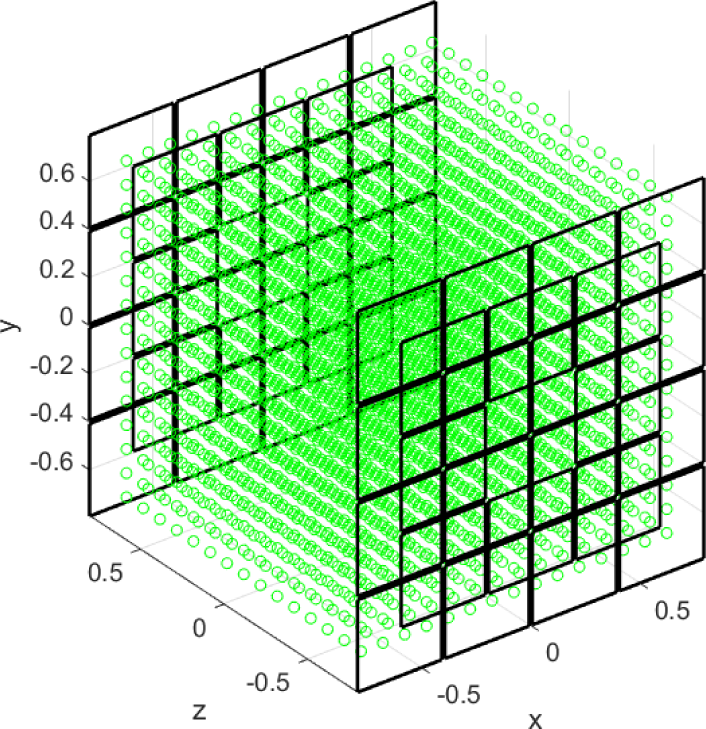
Coil coordinate system and grid of points (green) over which field noise was estimated.

Figures S9-S12 show maps of the spatial variation of the magnetic field over the volume between the coil panels. The noise at the centre of the coil planes is <20 fT/√Hz in each direction which is comparable to the OPM sensitivity. The strength of the magnetic field noise increases as proximity to the coils increases.

**Figure S9:**
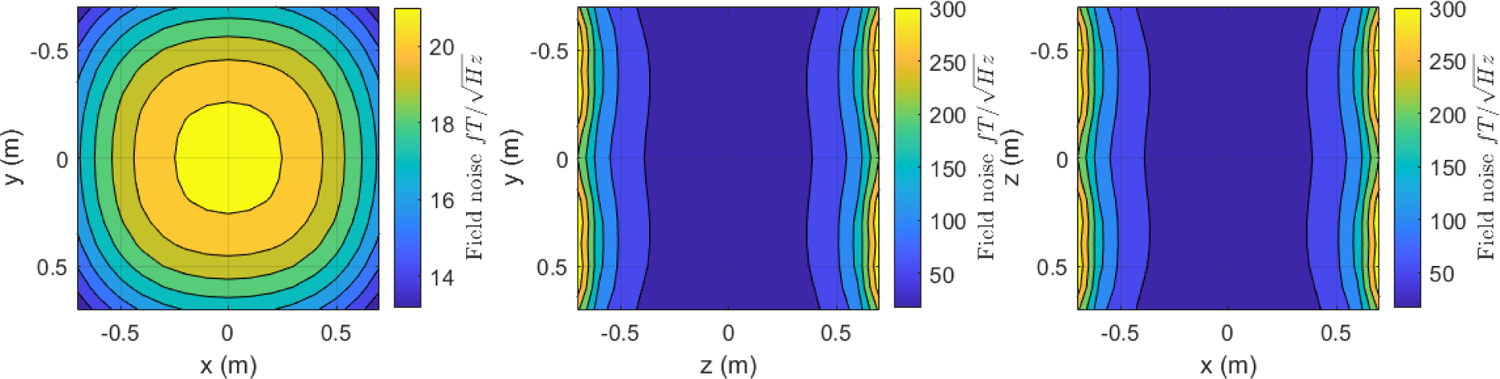
Map of the noise estimate of |B| in fT/√Hz. Slice 1 shows variation in the x − y plane at z = 0. Slice 2 shows variation in the x − y plane at x = 0. Slice 3 shows variation in the x − z plane at y = 0.

**Figure S10:**
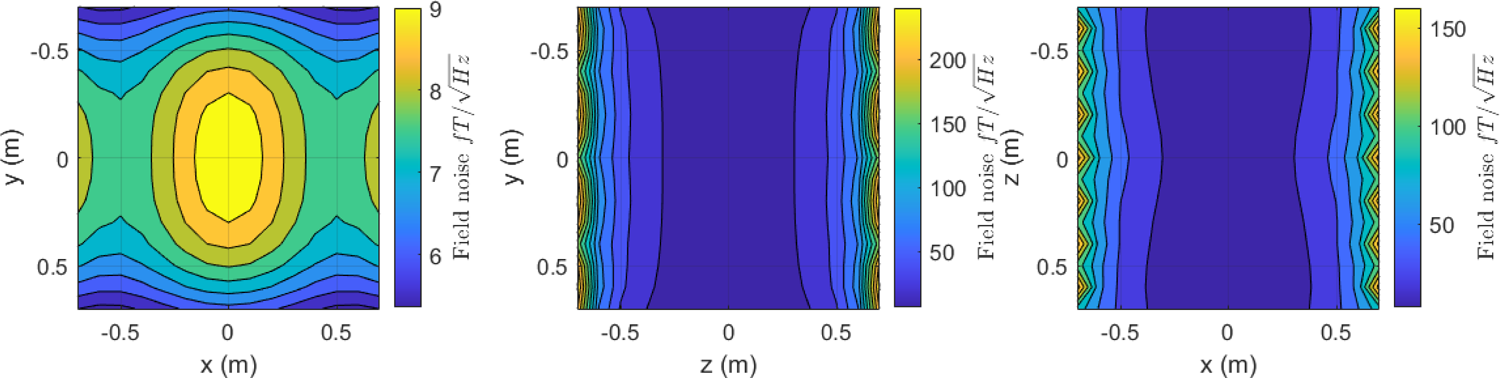
Map of the noise estimate of B_x_.

**Figure S11:**
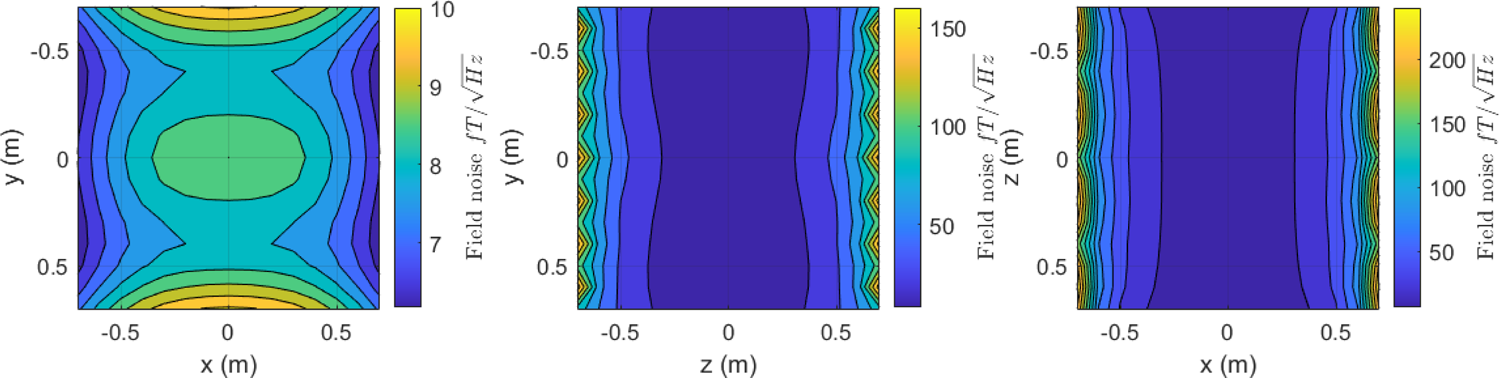
Map of the noise estimate of B_y_.

**Figure S12:**
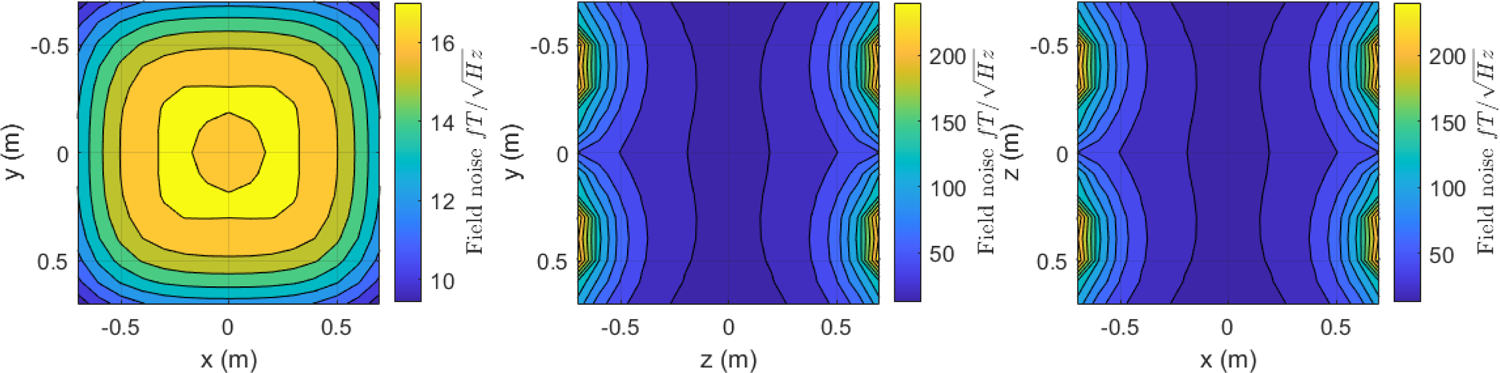
Map of the noise estimate of B_z_.

